# Direct Photon-by-photon Analysis of Time-Resolved Pulsed Excitation Data using Bayesian Nonparametrics

**DOI:** 10.1101/2020.07.20.212688

**Authors:** Meysam Tavakoli, Sina Jazani, Ioannis Sgouralis, Wooseok Heo, Kunihiko Ishii, Tahei Tahara, Steve Pressé

## Abstract

Lifetimes of chemical species are typically estimated, across each illuminated spot of a sample, by either fitting time correlated single photon counting (TCSPC) decay histograms or, more recently, through phasor analysis from time-resolved photon arrivals. While both methods yield lifetimes in a computationally efficient manner, the performance of both methods is limited by the choices made when fitting a TCSPC histogram. In addition, phasor analysis also requires setting the number of chemical species by hand before lifetimes can be determined. Yet the number of species itself is encoded in the photon arrival times collected for each illuminated spot and need not be set by hand *a priori*. Here we propose a direct photo-by-photon analysis of data drawn from pulsed excitation experiments to infer, simultaneously and self-consistently, the number of species and their associated lifetimes from as little as a few thousand photons for two species. We do so by leveraging new mathematical tools within the Bayesian nonparametric (BNP) paradigm that we have previously exploited in the analysis of single photon arrivals from single spot confocal microscopy. We benchmark our method on simulated as well as experimental data for one, two, three, and four species with data sets from both immobilized and freely diffusing molecules at the level of one illuminated spot.

**SUMMARY:** Photon arrivals obtained from fluorescence experiments encode not only the lifetimes of chemical species but also the number of chemical species involved in the experiment. Traditional methods of analysis, such as phasor methods and methods relying on maximum likelihood or (parametric) Bayesian analysis of photon arrivals or photon arrival histograms of TCSPC data, must first ascertain the number of chemical species separately and, once specified, determine their associated lifetimes. Here we develop a method to learn the number of fluorescence species and their associated lifetimes simultaneously. We achieve this by exploiting Bayesian nonparametrics. We benchmark our approach on both simulated and experimental data for one species and mixtures of two to four species.

## INTRODUCTION

Fluorescence microscopy provides a means to selectively monitor the dynamics and chemical properties of fluorophores or labeled molecules.^1–13^ In this study, our focus is on methods that use pulsed illumination^14–18^ or illumination modulated at a fixed frequency^18–23^ at one spot. Photon arrival times assessed in these methods encode critical information on the excited state lifetime or the number of different chemical species contained in the sample under imaging. This is the basis of lifetime imaging^13,24–27^ that has been used to reveal information on local pH,^28,29^ oxygenation^28^ and other cellular metabolic traits^23,30^ reporting back on the breadth of cellular microenvironments.

Maximum likelihood or traditional (parametric) Bayesian methods^31–35^ are common starting points in the analysis of photon arrivals or photon arrival histograms derived from pulsed illumination, *i.e*., time-correlated single photon counting (TCSPC) data.^2,36–38^

In pulsed illumination,^39,40^ photon arrival times are analyzed,^41–44^ under the assumption of a known number of molecular species with unknown lifetimes to be determined.^31–35,45–50^ This approach is best illustrated in discussing photon arrival histograms which are typically fitted using multi-exponentials^49,51^ to identify the lifetime of each species. That is, lifetimes, *τ_m_*, and the weights of the *m^th^* lifetime component, *a_m_*, are modeled and determined using multi-exponential decay fits of the form

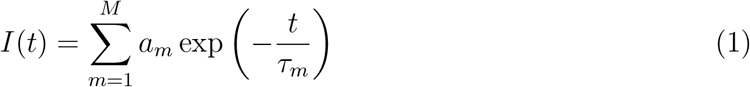

where *I*(*t*) is the intensity of photons arriving at time *t*.

In Eq. 1, the number of exponential components, *M*, must be specified before the data can be used to find *τ*_1_, ⋯, *τ_M_* and *a*_1_, ⋯, *a_M_*. Typically, *M* is specified according to some goodness-of-fit metric that safeguards against over-fitting^33^ as we discuss in the Supplementary Information. Indeed, within a maximum likelihood or parametric Bayesian paradigm, too large an *M* must be penalized according to *post hoc* criteria. ^52–55^ Other methods for deducing *M* rely on pole decompositions^56^ or Laplace-Padé expansions^57^ requiring exceedingly large data sets.

Another general method of analysis of lifetime data relies on phasors.^58–62^ Phasor analysis is appropriate for data from samples illuminated by light whose intensity is modulated at a fixed frequency.^21,58,63–65^ In this case, the intensity of the light emitted by the sample is also modulated and phase shifted.^18,59^ In particular, for a modulation frequency of *ω*, the measurements may be used to obtain the phase shift, *ϕ*, and the intensity modulation ratio, *m* (see Fig. S9). The phase shift and intensity modulation ratio, in turn, determine two coordinates (*G, S*) in a “phasor plot”

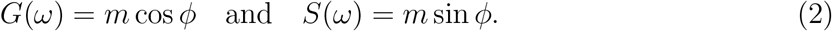

Lifetime values of the photon emitting fluorophores can then be deduced from the points on the phasor plot. ^60–62^

Phasor analysis is especially intuitive as it allows us to immediately deduce whether more than one lifetime component is present. ^66,67^ In particular, mono-exponential lifetimes fall somewhere on the semicircle of radius 1/2 beginning at coordinate (1, 0) and moving counterclockwise to (0, 0); see Fig. 1. Deviations thereof imply a mixture of lifetimes. Full details are provided in the Supplementary Information. A variant of phasor analysis also holds for pulsed excitation. ^60,68,69^ The advantages and drawbacks of phasors analysis are similar to those of the direct analysis of photon arrivals or histograms of photon arrivals from TCSPC data in that the number of species must be known in advance. What is more, the retrieval of lifetime information from phasor analysis requires independent knowledge of not only the number of species but, often, it also requires the lifetimes of all but one unknown species whose lifetime is to be determined from a mixture of chemical species; ^27,60,70,71^ see Fig. 1.

**Figure 1:**
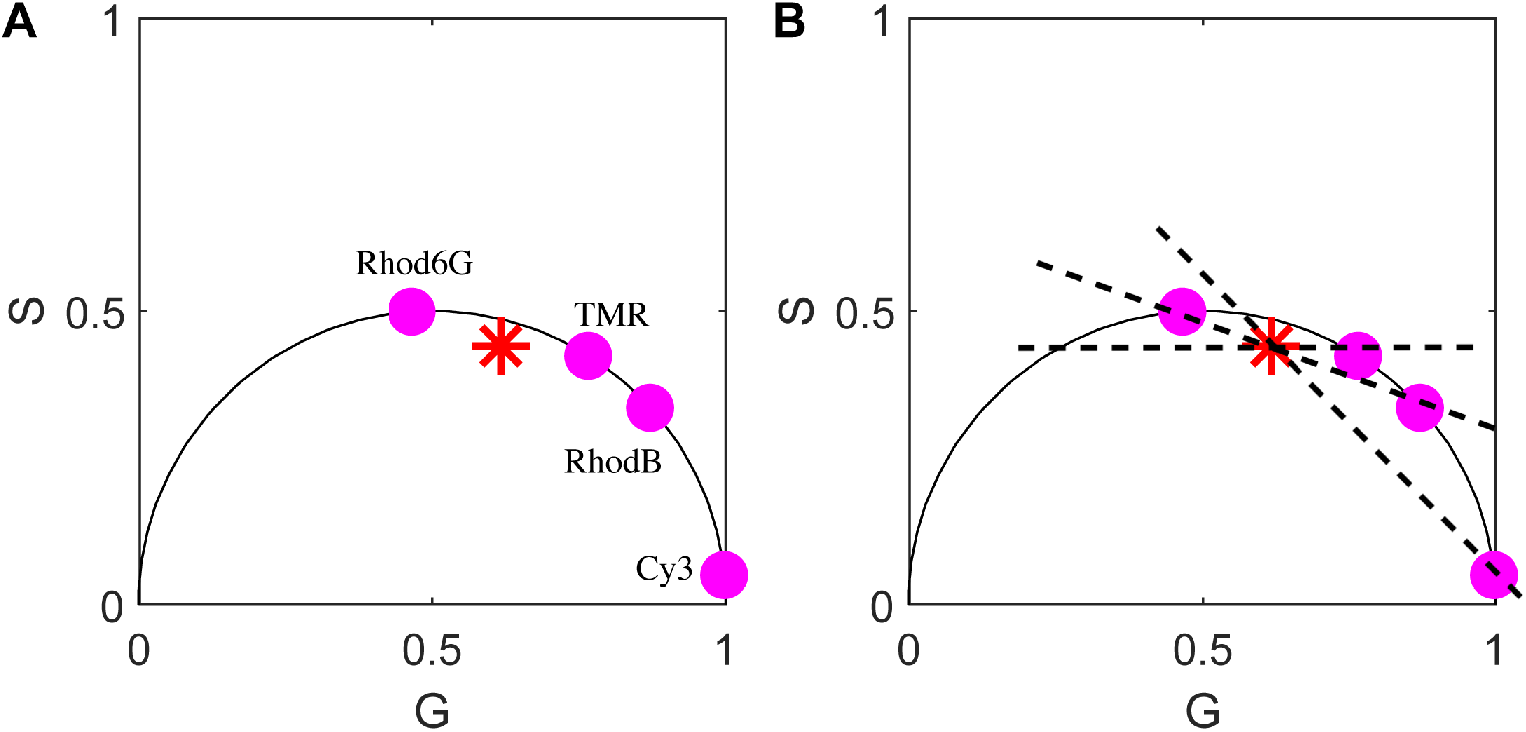
Phasor analysis provides the lifetimes of chemical species but not an independent measure of the number of chemical species. In panel (A) appears a typical phasor plot as expected when a mixture of 4-component mixture, red star, (Rhod-6G, TMR, RhodB, and Cy3) is subject to pulsed illumination. From this figure, it is not possible to discern the number of chemical species contributing to the phasor plot. What is more, as we can see in the panel (B), if we assume 2 species, many choices of lifetimes could be warranted by the data as evidenced by the placement of the dashed diagonal lines. The point of intersection of these diagonal lines with the phasor plot’s hemisphere would be needed to deduce the lifetimes of a 2-component mixture if we had hypothesized this mixture to be composed of 2 species (as opposed to the correct number, 4). The panel (B) superposes the phasor plots for each species measured independently. Their mixture is what yields the subfigure on the left whose identity as a 4-component mixture is not apparent.

While both approaches we have just described, direct photon analysis and phasors, yield lifetimes in a computationally efficient manner, their greatest limitation is the requirement that the number of species, *M*, be pre-specified as it otherwise *cannot* be learned independently although, in principle, it is encoded in the data. Yet, learning the number of species is critical as it may be unknown prior to collecting data for a number of reasons.^68,72–74^ At higher computational cost, we could learn not only the number of species, but even full joint distributions over the possible number of species as well as their associated lifetimes which are encoded in the photon arrivals. That is, we could determine the relative probability over having 3 versus 4 species, say, not just the most probable number of species. Ideally, to allow for higher flexibility in the experimental setting, we need to achieve this with the same or fewer photon arrivals than is required in direct photon and phasor analysis to reveal the lifetimes alone. In order to do so, we need to relinquish the traditional (parametric) Bayesian paradigm that assumes a fixed model structure (*i.e*., a fixed number of species).

We have previously exploited the Bayesian nonparametric (BNP) paradigm^75–78^ to analyze single photon arrival time traces in order to learn diffusion coefficients from minimal photon numbers drawn from single spot confocal experiments.^10,79^ Traditionally, such photon arrivals were analyzed using tools from fluorescence correlation spectroscopy where very long traces were collected and auto-correlated in time. Just as with the problem at hand, the direct photon-by-photon analysis demanded a different approach as the stochastic number of molecules contributing photons was unknown and an estimate of that number deeply impacted our diffusion coefficient estimate. It is for this reason that we invoked the non-parametric paradigm there. In particular, the BNP paradigm is also preferred here on this basis: assuming an incorrect number of species, when these and their associated lifetimes are assumed unknown, leads to incorrect lifetime estimates for each species; see Fig. 2. This further begs the question as to whether fits of the data with different, incorrect, models can be compared in the first place.

**Figure 2:**
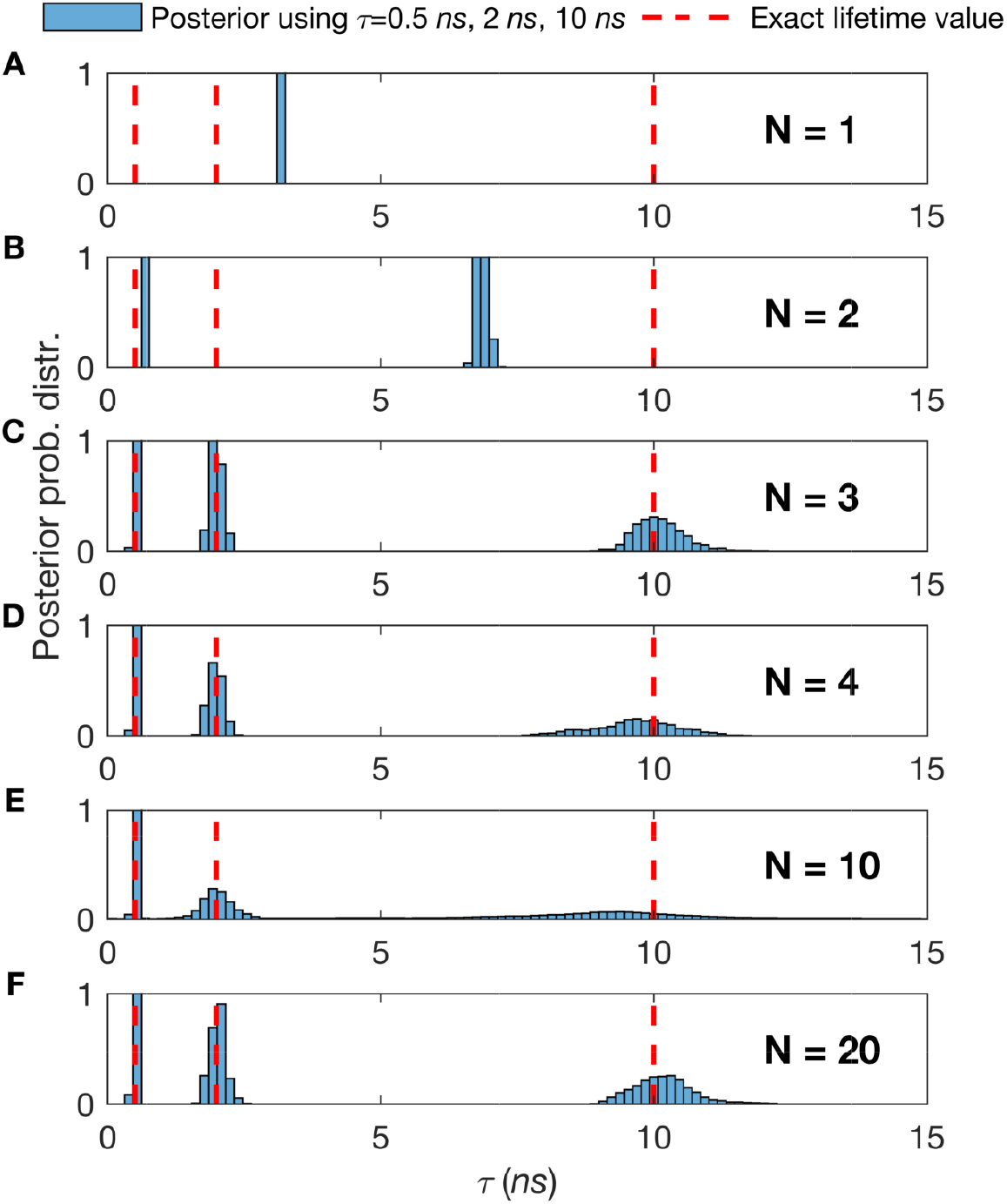
The number of species assumed in the analysis directly impacts the lifetimes ascribed to those species. Thus, we need an independent method to estimate species numbers. (A-F) We generate synthetic traces with three species with a total of 2 × 10^4^ photon arrivals and lifetimes, *τ*, of 0.5 *ns*, 2 *ns*, and 10 *ns*. To estimate the *τ* within the normal (**i.e**., parametric) Bayesian paradigm, we start by assuming the following number of species, *N* =1 (A), *N* = 2 (B), *N* = 3 (C), *N* = 4 (D), …, *N* =10 (E), …, and *N* = 20 (F). The good fit provided by *N* > 2 and the mismatch in the peak of the posterior distribution over the lifetime and correct value of the lifetime (red dotted line) in all others underscores why it will become critical for us, or any method analyzing single photon data in the context of confocal microscope experiments, to correctly estimate the number of species contributing to the trace in order to deduce chemical parameters such as lifetime.

Here we propose a method that exploits BNPs^80^ to learn species and their associated lifetimes with as few photons as possible using pulsed illumination from a single illuminated spot. As with any inverse methods, in BNPs we start from the data: namely the time lag between the peak of the pulse and the detection time of the photon termed “microtime” discussed in more detail later on. To be precise, each species is defined as contributing photons some time after pulsing dictated by an exponential distribution with a decay constant (lifetime) unique to that species. Just as we treat model parameters as random variables in the parametric Bayesian paradigm, within the BNP paradigm, we treat models themselves as the random variables and try to learn full posterior distributions over the number of species.

The advantages of using BNPs are four-fold: 1) we can learn full posterior distributions over species present in the measurements which is especially relevant for datasets with limited photons as the number of species becomes highly uncertain; 2) by resolving lifetimes and species from the raw photon arrivals directly, by contrast to processed data which necessarily contains less information, we can minimize photo-damage; 3) as a corollary to the previous point, we can monitor processes out-of-equilibrium where only few photons are available before chemical conversion into another species; 4) given long traces, we can exploit the additional data, if need be, to discriminate between species with small differences in lifetimes.

## RESULTS

Our goal is to characterize quantities that describe molecular chemistry at the data-acquisition timescales of TCSPC with a focus on obtaining lifetime estimates and the number of chemical species. In order to estimate lifetimes, we also estimate intermediate quantities, such as the fraction of different species contributing photons as detailed in the method section.

Within the BNP approach,^81–83^ our estimates take the form of posterior probability distributions over unknown quantities. These distributions combine parameter values, probabilistic relations among different parameters, as well as the associated uncertainties. To quantify this uncertainty, we calculate a posterior variance and use this variance to construct error-bars (*i.e*., credible intervals). As follows from Bayesian logic, the sharper the posterior, the more conclusive (and certain) the estimate. ^79,81,84^

### Method Validation using Synthetic Data

To show the robustness of our method, we generate synthetic traces for immobilized molecules with: i) variable data set sizes, Fig. 3 involving multiple species, Fig. 4; ii) variable fraction of molecules contributing photons from different species, Fig. 5; and iii) variable difference of lifetimes for mixtures of lifetimes, Fig. 6. All parameters not explicitly varied are held constant across all figures. The parameters not varied are held fixed at the following baseline values: lifetime between 1 ns and 10 ns which is the typical lifetime range of a fluorophore, ^18,85^ two species which is most frequent in related studies, ^18,19,23^ and fraction of molecules contributing photons from different species set at 50% : 50%.

**Figure 3:**
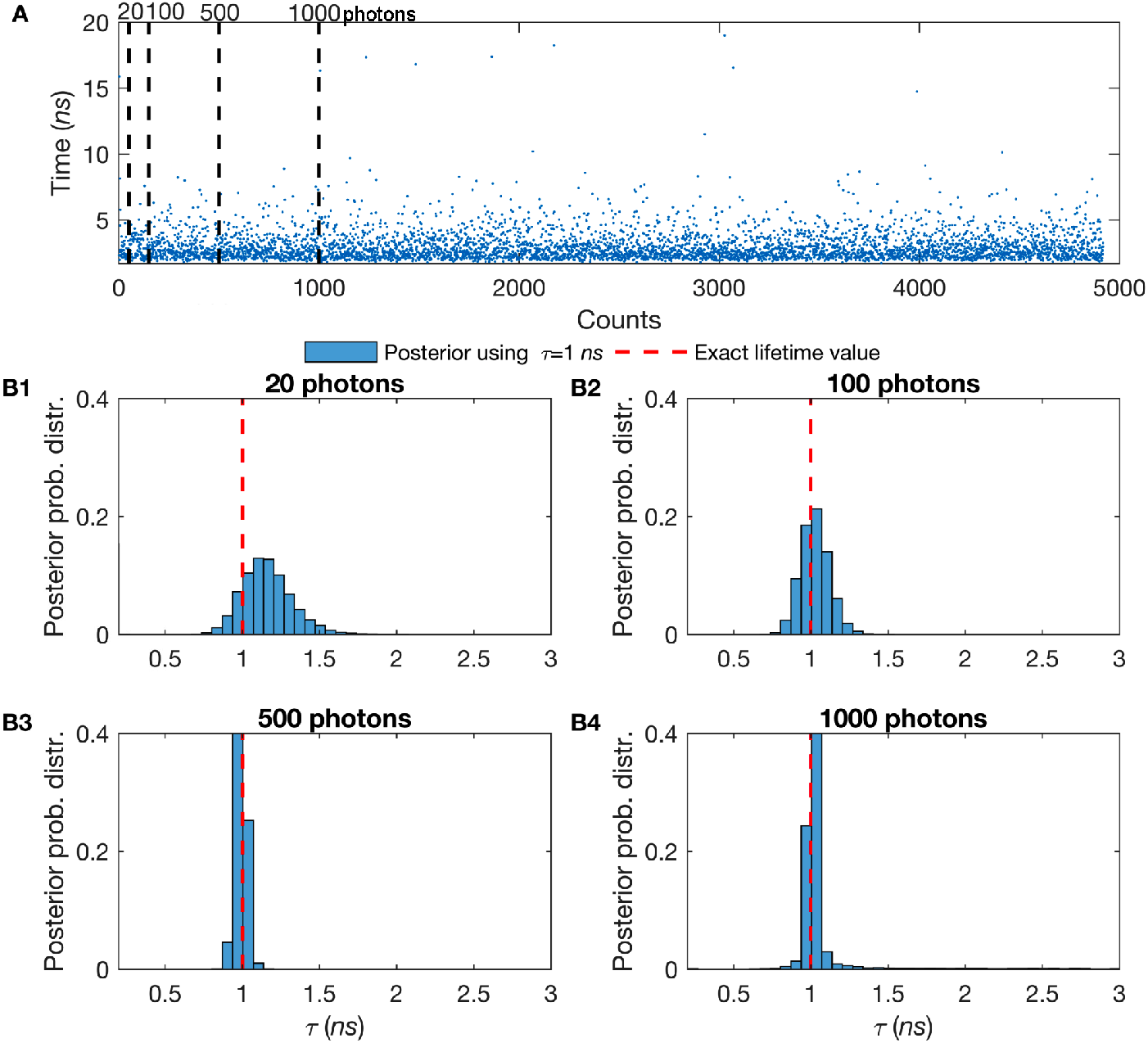
The greater the number of detected photons, the sharper the molecular lifetime estimate. (A) Here, we work on single species lifetime while all molecules are immobilized. The synthetic trace generated using a lifetime of *τ* = 1 *ns*. The blue dots represent single photon arrival times (y-axis) recorded after each excitation pulse (x-axis). We consider the excitation pulse as a Gaussian IRF (Eq. 4) occurs at a frequency of 40 *MHz* with standard deviation of 0.1 *ns*. (B1) In the analysis to determine lifetimes, we first start with just 50 photons, first black-dashed line in panel (A), and gradually increase the number of photons considered in the analysis to (B2) 100, second black-dashed line in panel (A), (B3) 500, third black-dashed line in panel (A), and (B4) 1000 photons, last black-dashed line in panel (A). The ground truth for the lifetime is known (as this is synthetic data) and it is shown by the red-dashed line.

**Figure 4:**
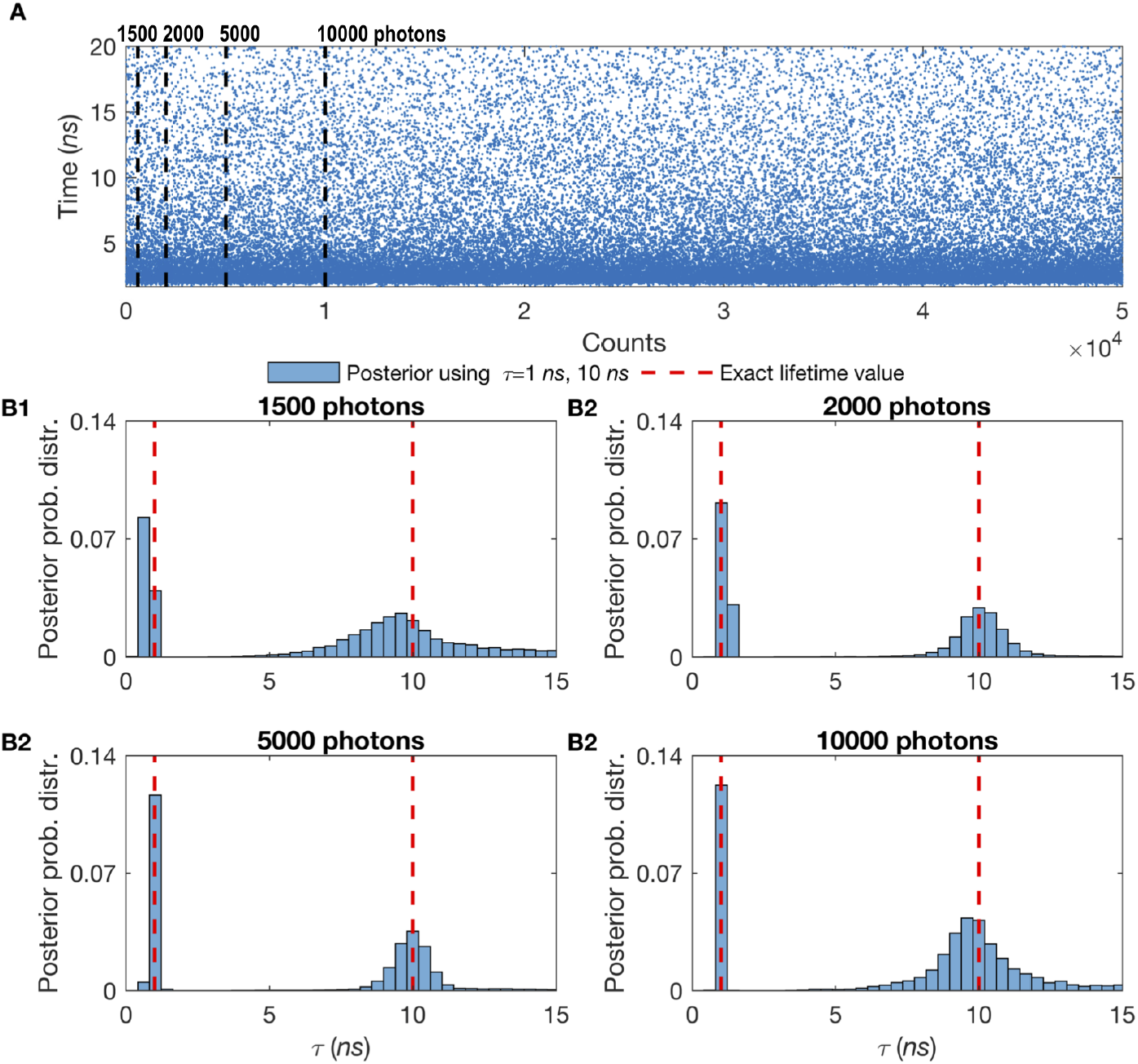
Effect of the number of detected photons on two simultaneous molecular lifetime estimates. The more photons collected, the sharper the lifetime estimate for the case of two species. (A) Here, we use mixtures of two species with different lifetimes while all molecules are immobilized. The synthetic data is generated using *τ* = 1 *ns* for the first species and *τ* = 10 *ns* for the second with equal ratio of molecules of each species (50% – 50%). The blue dots represent single photon arrival times detected after each excitation pulse. (B1) In the analysis to determine both lifetimes, we first start with just 1500 photons, first dashed line in panel (A), and gradually increase the number of photons to 2000 (B2), 5000 (B3), and 10000 (B4) photons. Here, all other features such as the frequency of acquisition and width of pulse are the same as in Fig. 3. Also, we follow the same red-dashed line convention as in 3. To see the results for more than two species see the Supplementary Information, Figs. S4 and S5.

**Figure 5:**
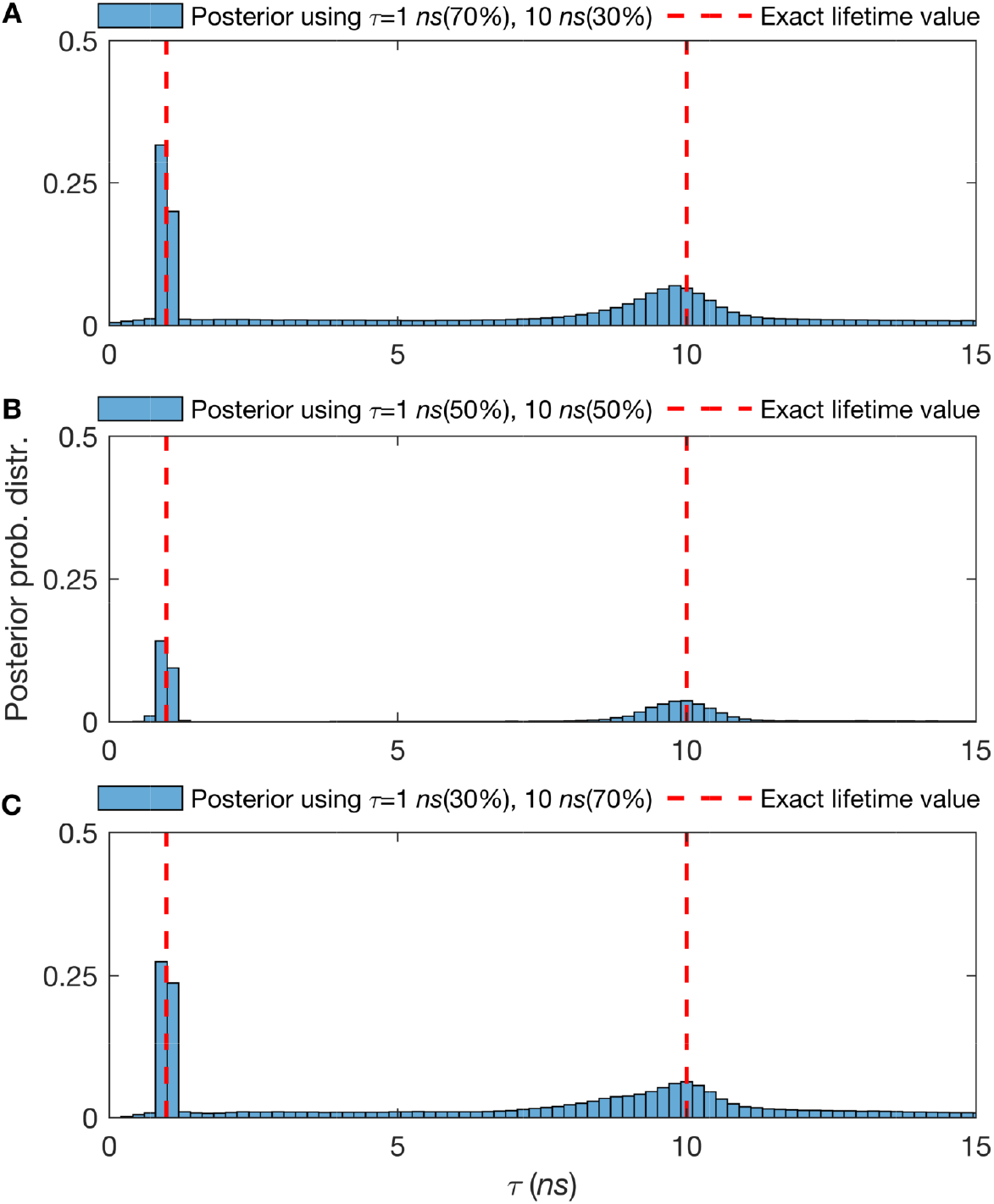
Effect of the relative fraction of contributing molecules from different species on molecular lifetime estimates. Higher molecular contributions provide more photons per unit time and thus sharper lifetimes estimates. (A-C) The posterior probability distributions of traces with lifetimes of 1 *ns* and 10 *ns*, with 3000 total photons and fraction of contributing molecules from different species of 70% – 30%, 50% – 50% and 30% – 70% respectively. Here, all other features such as the frequency of acquisition and width of pulse are the same as in Fig. 3. Also, we follow the same red-dashed line convention as in 3. For more details see Supplementary Information Fig. S6.

**Figure 6:**
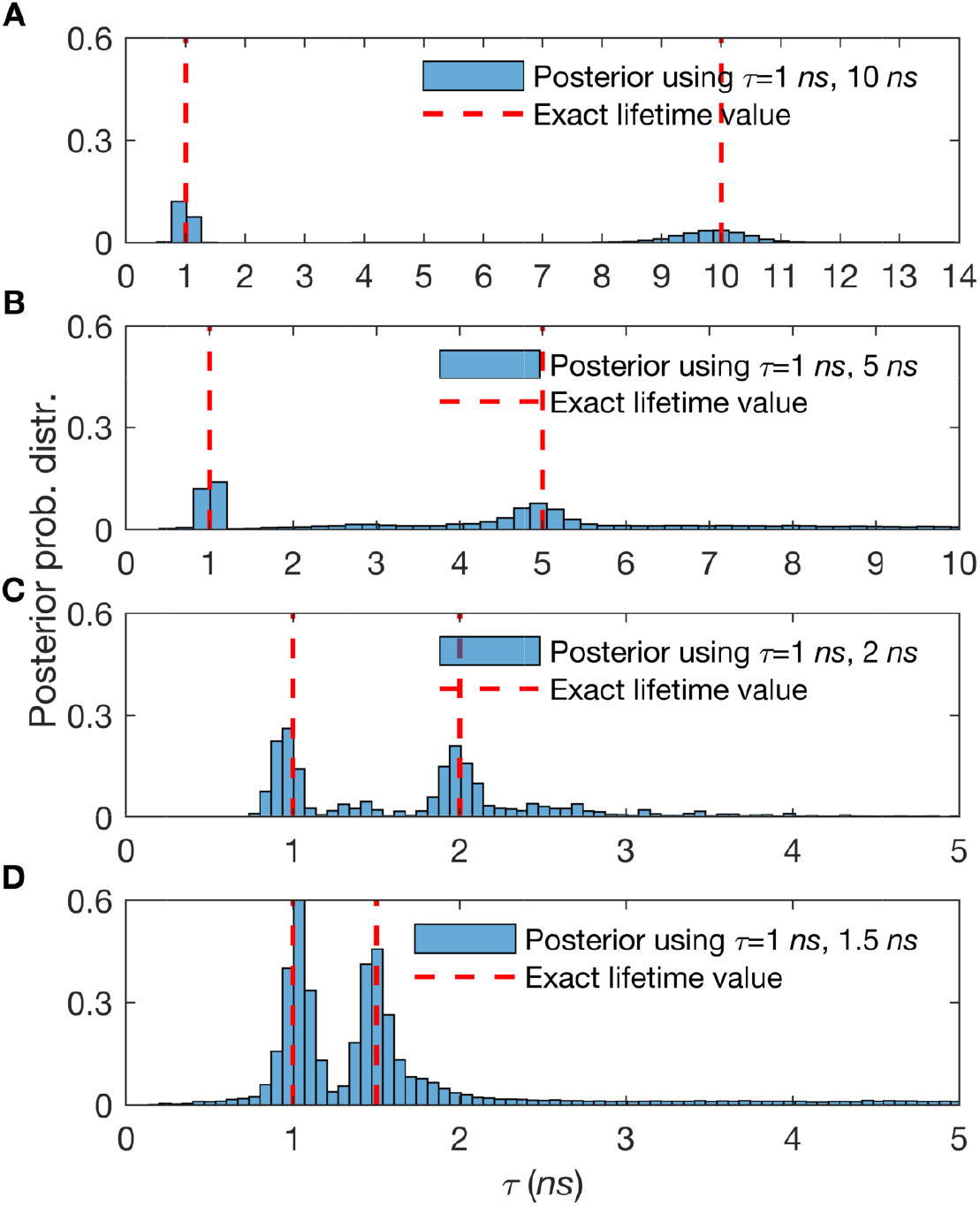
Lifetime resolution for double species lifetimes. Here we show posterior probability distributions over estimated lifetimes. In The synthetic traces acquired contain 3000 to 20000 photon arrivals and start in (A) with well separated lifetimes of 1 *ns* and 10 *ns* (≈ 3000 photons) before gradually considering less well separated lifetimes such as in (B) where the lifetimes are 1 *ns* and 5 *ns* (≈ 3000 photons), in (C) where the lifetimes are 1 ns and 2 ns (≈ 10000 photons), and in (D) where the lifetimes are 1 ns and at last 1.5 ns (≈ 20000 photons). The fraction of molecules contributing photons from different species is even split (50% – 50%). Here, all other features, such as the frequency of acquisition and width of pulse, are the same as in Fig. 3. Also, we follow the same red-dashed line convention as in 3.

Also, in the Supplementary Information, we worked with cases involving three and four different species (as opposed to a just one or even two species) as this scenario presents the greatest analysis challenge because very few photons, and thus little information, is gathered on each species. In a similar spirit, we also default to short traces that highlight the value of analyzing data in its rawest form. As the mathematics remain unaffected, and this scenario reflects the reality of many experiments, we show in the Supplementary Information, Figs. S1 and S2, results for freely diffusive molecules.

#### Number of photons

We benchmark the robustness of our approach with respect to the length of the trace (*i.e*., the total number of photon arrivals) at fixed number of species, lifetime, and molecule photon emission rate. For instance, to obtain an estimate of the lifetime within 10% of the correct result in the one species case, our method requires only ≈ 100 photons (emitted from the species of interest). In the case of two species, our proposed BNP approach requires only ≈ 3000 photons; see Figs. 3 and 4. To determine how many photons were required by our method, we chose the mean value of the lifetime posterior, and measure the percentage difference of this mean to the ground truth known for these synthetic traces.

In general, the numbers of photons demanded by our method are minimal though the absolute number depends on a broad range of experimental parameter settings. This is the reason why, throughout this work, we explore different settings—holding all other settings fixed—in subsequent subsections as well as the Supplementary Information.

Another important concept, illustrated in Figs. 3, and 4 that will keep re-appearing in subsequent sections, is the concept of a photon as a *unit of information*. The more photons we have, the sharper our lifetime estimates. This is true, as we see in these figures, for increasing trace length. Similarly, as we will see in subsequent subsections, we also collect more photons as we increase the contribution of labeled molecules (and thus the number of molecules contributing photons to the trace).

#### Mixtures of different species contributing photons

To test the robustness of our method when different species contribute an uneven number of photons, we simulate data with 70% of the population in species 1 and 30% in species 2 (Fig. 5A). We also considered fractions of contributing molecules from different species of 50% : 50% (Fig. 5 B), and 30% : 70% (Fig. 5 C). For all cases, the lifetimes were fixed at 1 ns and 10 ns for ≈ 3000 photon arrivals. Fig. 5 summarizes our results and suggests that posteriors over lifetimes are broader—and thus the accuracy with which we can pinpoint the lifetimes drops—when the contribution of labeled molecules is lower. Intuitively, we expect this result as fewer species within the confocal volume provide fewer photons and each photon carries with it information that helps refine our estimated lifetimes.

#### Lifetime resolution

We repeat the simulations with two species and ask about how many photons are required to resolve similar lifetimes. Here, we have presented the dependency of the time resolution to the number of collected photons in Fig. 6. As expected, the number of photons required to resolve increasingly similar lifetimes grows as the ratio of lifetimes approaches unity. However, this also suggests that if we were to resolve species of similar lifetimes, we could use the amount of data typically used in TCSPC or phasor analysis to resolve these while TCSPC or phasor analysis would still require an additional order of magnitude more data. As we noted earlier, both TCSPS and phasor analysis must impose by hand the number of species while, in our method, the number of species are learnt. Moreover, if we know number of species we require even fewer photons that we mentioned earlier.

### Estimation of physical parameters from experimental data

To evaluate our approach on real data, we used experimental data collected under a broad range of conditions. That is, we used measurements from different fluorophores, namely Cy3, TMR, Rhod-B, and Rhod-6G. The lifetimes for these dyes are first benchmarked by fitting TCSPC photon arrival histograms from entire traces and comparing with published-values. ^86–89^

Figs. 7, 8 and 9 were collected using the Rhod-B and Rhod-6G dyes and these results were used to benchmark the robustness of our method on individual species as well as mixtures of species with a variable fraction of chemical species contributing photons. In the Supplementary Information, Fig. S7, we show more experimental results for cases involving more than two species.

**Figure 7:**
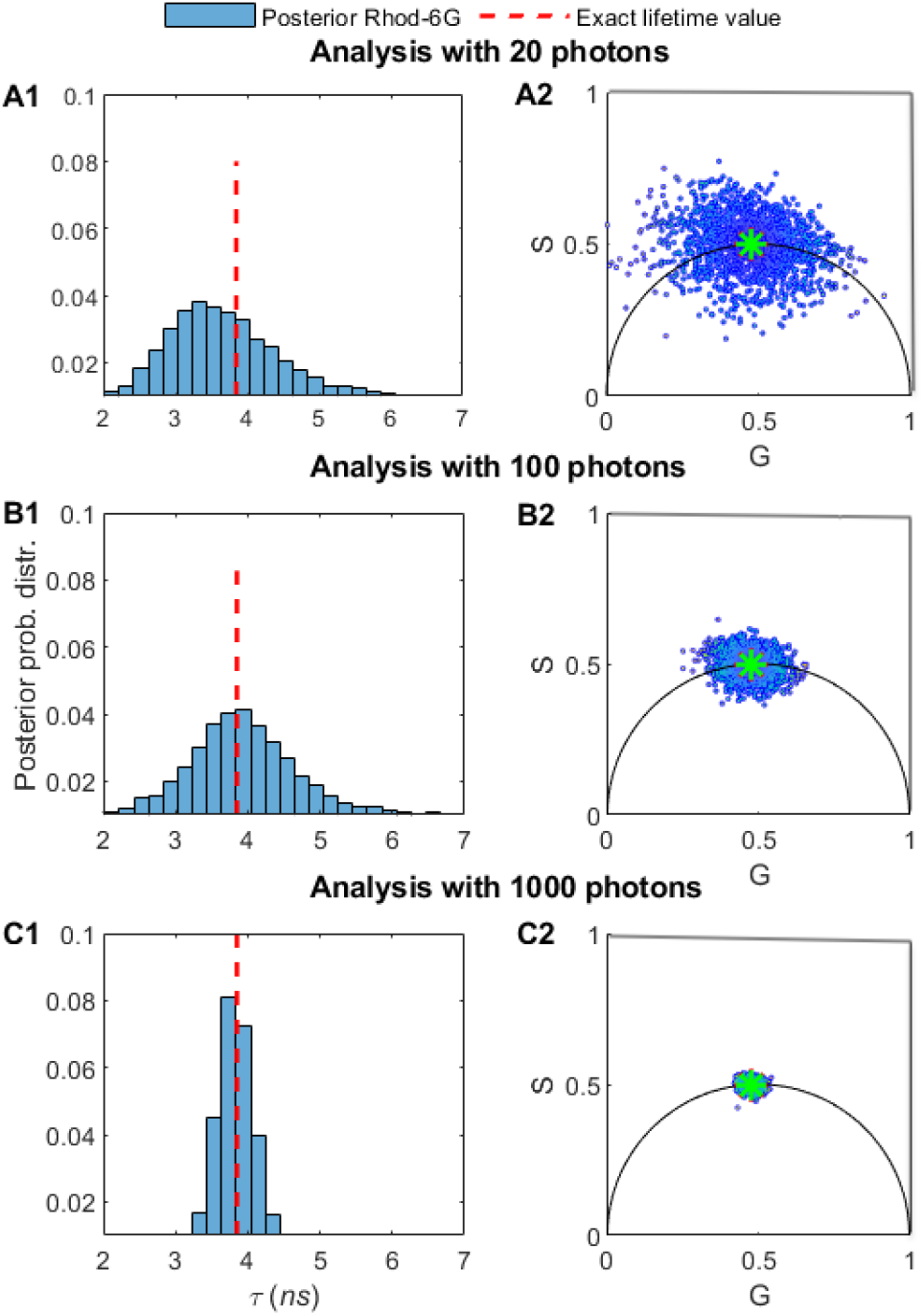
Comparison of the number of photons needed to assess the lifetimes of Rhod-6G. (A1) We use 20 photons from experimental time trace Rhod-6G. For visualization purposes only, we show the corresponding phasor plots in (A2). In (B1-B2) and (C1-C2) we repeat the analysis for 100 and then 1000 photons. Using our method relying on BNPs, the estimated lifetimes are: (A1) *τ* = 3.10 *ns*, (B1) *τ* = 3.95 *ns*, and (C1) *τ* = 3.91 *ns*. The excitation pulses occur at frequency of 40*MHz* and we assume a Gaussian shape with standard deviation of 0.1*ns*. The ground truth (red-dashed lines) is obtained using TCSPC photon arrival histogram fitting when analyzing the whole time trace. In our BNP analysis, we do not pre-specify the number of species, we learn them alongside the associated lifetimes.

**Figure 8:**
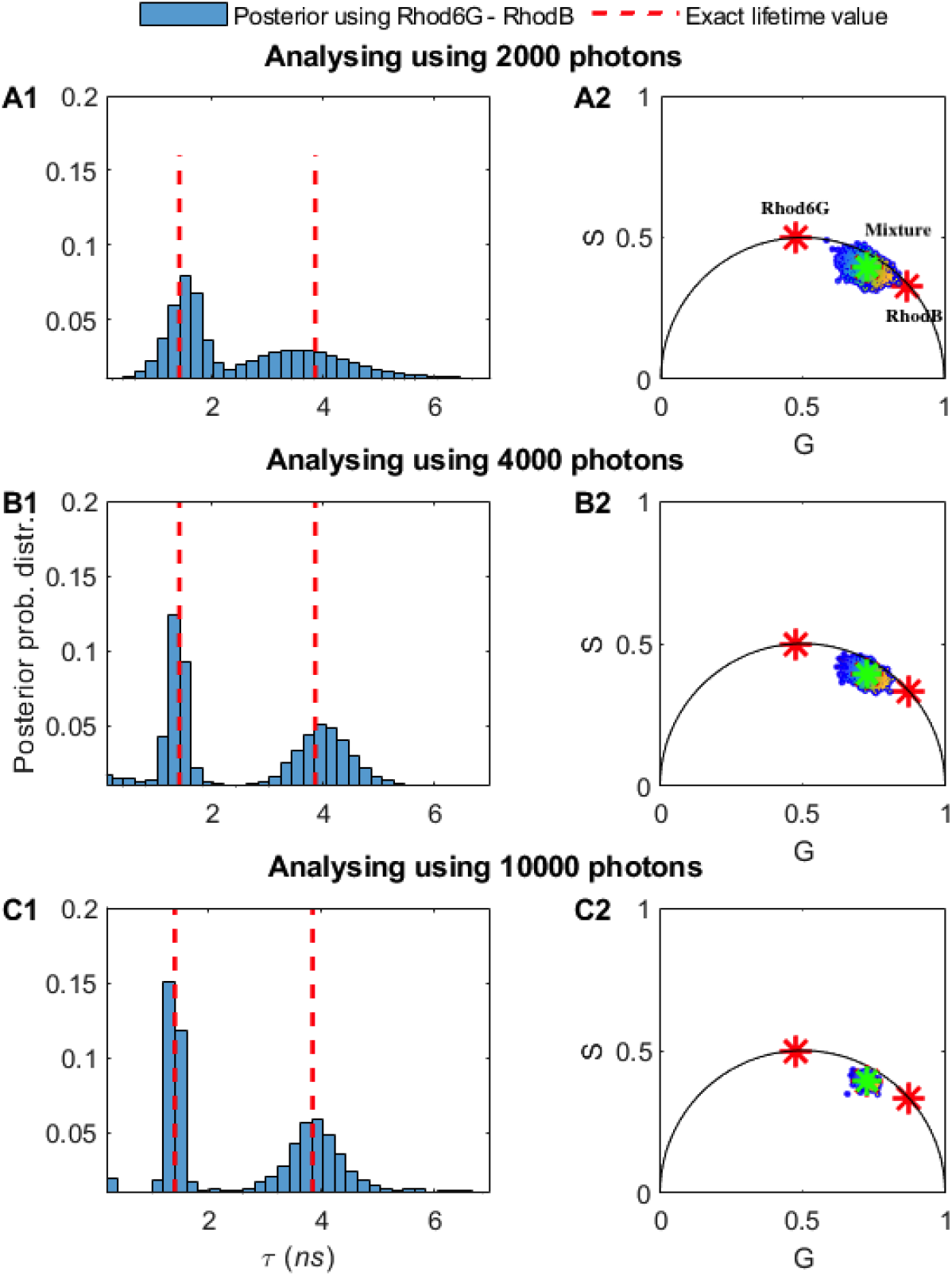
Comparison of the number of photons needed to assess the lifetimes of mixtures of Rhod-B and Rhod-6G. In (A1) we use 2000 photons. For visualization purposes only, we show the corresponding phasor plots in (B1). In (B1-B2) and (C1-C2) we repeat the analysis for 4000 and then 10^4^ photons. Using BNPs, the estimated lifetimes: (A1) *τ* = 1.44, 3.39 *ns*, (B1) *τ* = 1.42, 3.96 *ns*, and (C1) *τ* = 1.41, 3.90 *ns*. Here all other features such as the ground truth (red-dashed lines), frequency of acquisition, etc. are the same as in Fig. 7. The green star on (A2)-(C2) is the location of mixture of 2 species when we use whole trace and the red stars show the location of the single species lifetime, for visualization purposes only, whose lifetimes we independently know from experiments on individual species.

**Figure 9:**
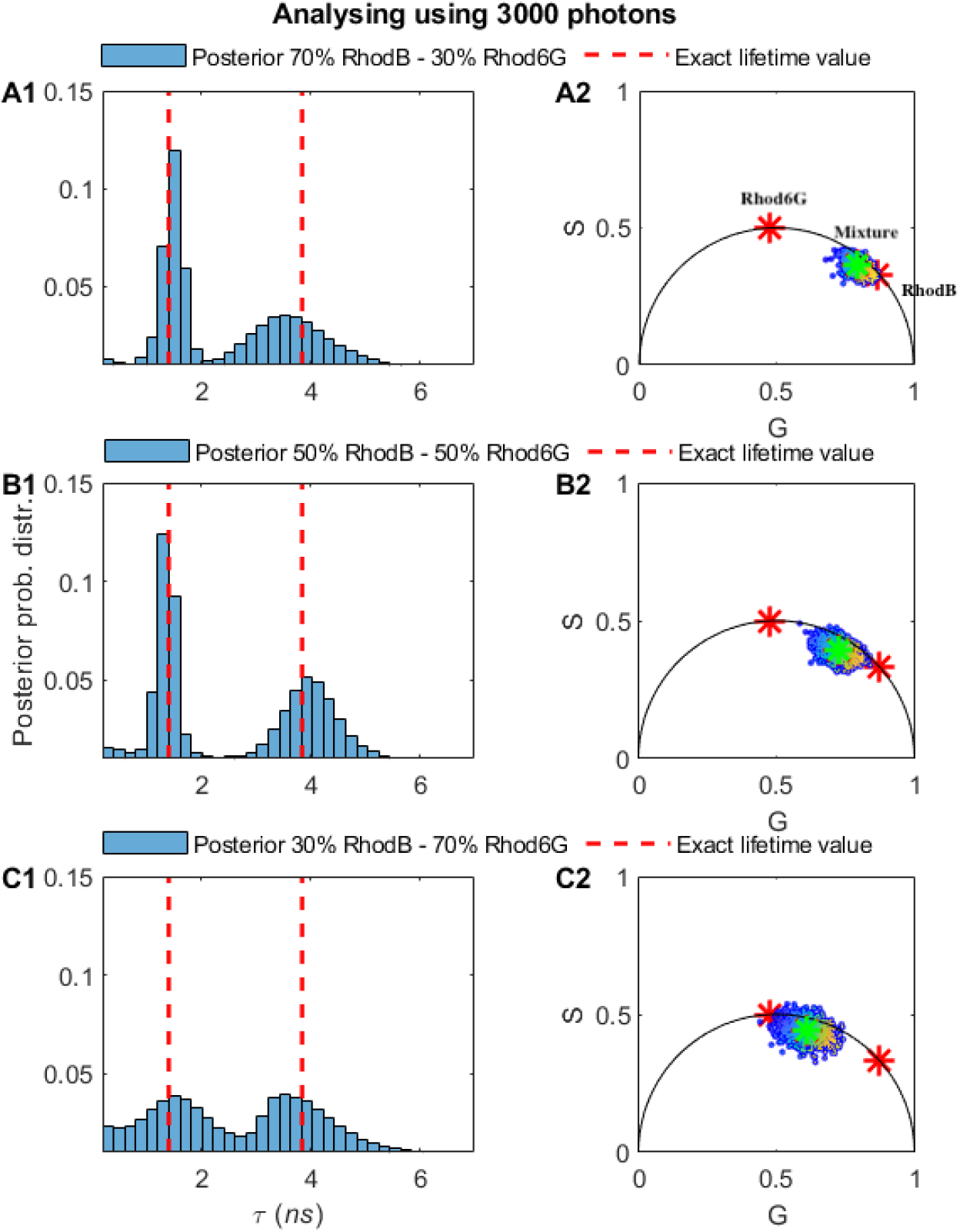
Effect of the fraction of molecules contributing photons from different species on molecular lifetime estimates. Higher molecular contributions provide more photons per unit time and thus sharper lifetime estimates. The experimental trace is selected using two species, Rhod-B and Rhod-6G, with a total of ≈ 3000 photon arrivals with a different fraction of photons derived from different species (70% – 30%)(A1), 50% – 50% (B1), and 30% – 70% (C1). The estimated lifetimes using BNPs are: (A1) *τ* = 1.44, 3.39 *ns*, (B1) *τ* = 1.42, 3.96 *ns*, and (C1) *τ* = 1.41, 3.90 *ns*. Here all other features such as the ground truth (red-dashed lines), frequency of acquisition, etc. are the same as in Fig. 7. The green and red stars on subfigures (A2)-(C2) are explained in the caption of Fig. 8.

In Fig. 7, we verified our method on Rhod-6G with respect to the total number of photon arrivals. The first important conclusion is that we need ≈ 100 photons to obtain an estimate of the lifetime within 10% of the correct result (as obtained from our benchmark). For two or more species, the situation for phasor analysis, TCSPC photon arrival histogram fitting, or direct analysis of photon arrivals using parametric Bayesian methods or maximum likelihood grows more challenging. The number of species cannot be independently determined and, assuming an incorrect number of species leads to incorrect lifetime estimates; see Fig. 1 for phasors and Fig. 2. Moreover, for all cases, we could reliably determine the ground truth (red-dashed lines) from TCSPC photon arrival histogram fitting when using the whole trace with all photons available. To be clear, we learn the number of species directly using BNPs and do not assume a number ahead of time.

Once more, the absolute number of photons demanded by our method depends on a broad range of experimental parameter settings. This is the reason, we explore different settings—holding all other settings fixed—just as we did with synthetic data in subsequent subsections as well as the Supplementary Information.

#### Benchmarking on experimental data using a different number of photons for mixtures of Rhod-B and Rhod-6G

Similarly to the synthetic data analysis appearing in Fig. 4, we benchmark the robustness of our approach with respect to the length of the trace (**i.e**., the total number of photon arrivals) given fixed lifetimes and fraction of chemical species contributing equal numbers of photons (50% : 50%). Again an important message here is that, for the values of parameters selected, we need ≈ 100 photons for single species, Fig. 7, and ≈ 3000 photons for double species, Figs. 8, and 9. For instance, to obtain an estimate of the lifetime to within 10% of the correct result for the case of two species, our method requires ≈ 3000 photons.

#### Benchmarking on experimental data using different fractions of Rhod-B and Rhod-6G

We start by evaluating our method on mixtures of Rhod-B and Rhod-6G but present in different amounts. Similarly to Fig. 5 for the analysis of two species from synthetic data, we show estimates of the lifetimes for two species, Rhod-B and Rhod-6G, present at 70% : 30% fraction (Fig. 9A), at 50% : 50% fraction (Fig. 9B), and at 30% : 70% fraction (Fig. 9C). Fig. 9 summarizes our results and suggests that posteriors over lifetimes are broader—and thus the accuracy with which we can pinpoint the lifetimes drops—when the contribution from the dye concentration for that species is lower. To obtain an estimate of the lifetime to within 10% of the correct result, our method requires ≈ 3000 photons directly emitted from the dye; for visualization purposes, the corresponding phasor plot is provided in Fig. 9. In the Supplementary Information, we show additional results for the case of three and four species which are additionally challenging for existing methods with different fractions of chemical species contributing photons.

## DISCUSSION

Across all spectroscopic and imaging applications, the photon is the basic unit of information.^79,90^ Decoding information directly from single photon arrivals, with as few photons as possible without binning or correlating or other pre-processing of the data, is the main focus of our data-centric analysis strategy. Yet decoding information directly from single photon arrivals presents fundamental model selection problems.

For example, in the case of FCS, if we are to learn diffusion coefficients directly from limited photon arrivals, we must know how to write down a likelihood or, put differently, we must know the number of molecules contributing photons that, in turn, dictate the form for the likelihood. ^79^ As we do not know how many molecules we have, and what the appropriate likelihood should be, we have a model selection problem. Similarly, for lifetime imaging, if we are to learn the lifetime of the chemical species contributing photons, we must also know the number of species in order to write down a conventional likelihood.

Traditional Bayesian methods do not have a direct solution to the model selection problem^80,82^ as they also require us to be able to write down a likelihood. That is, they consider a fixed model (and a fixed likelihood) and treat the model’s parameters as random variables of the posterior distribution. By contrast, BNP, which are a direct logical extension of parametric Bayesian methods, treat models alongside their parameters as random variables.^75,83,91–96^

This ability to treat models themselves as random variables is the key technical innovation that prompted the development of BNPs in the first place. BNPs make it possible to avoid the computationally infeasible task of first enumerating and second comparing all models for any associated parameter values to all other competing models and their associated parameter values.

The BNP approach to tackling lifetime image analysis that we propose here cannot replace phasor analysis^20,23,60,62,64,69,97^ or TCSPC photon arrival analysis under an assumed number of species^2,14,29,38,40,98^ for simple one component systems on account of their computational efficiency. However, at an acceptable computational cost, BNP approaches provide a powerful alternative. They give us the ability to: determine the number of species (and probabilities over them if the data are uncertain due to its sparsity or otherwise); use much less data to obtain lifetime estimates (and thus reduce photo-toxic damage to a light-sensitive sample); use longer photon arrival time traces, if available, to tease out small differences in lifetimes between species as BNP-based methods are more data efficient; probe processes resolved on faster timescales (again, as we require minimal photon numbers); exploit all information encoded in the photon arrivals (and thus not require separate control experiments, as often needed in phasor approaches, for the measurement of the lifetime of one species to determine the lifetime of a second species when a mixture of two species, say, is present).

As for the computational cost, obtaining lifetimes (to within 10% of the ground truth lifetime for a one-species for the parameters we used in Figs. 3 and 7 requiring ≈100 photons) takes 5 minutes on a typical scientific desktop as of the publication date of this paper (based on a system with 6G RAM, Core (TM) i7-2.67 GHz CPU). For a two-species mixture, Figs. 4 and 8, under the same parameters and requiring 3000 photons, it was a modest increase to 15 minutes. The point, here, is that the analysis of single or multi-species data *can* be performed with an average desktop computer and it does not necessarily require high performance computing facilities.

The real strength of BNP becomes clear when we reach two, three, four or possibly even more species. Beyond being able to work with low photon counts, another key advantage of our method is its flexibility. The ability to use BNP, and treat models as random variables, in lifetime imaging is the real point here and, as such, our framework can be adapted to treat a range of experimental setups.

In particular, our framework can straightforwardly be adapted to treat: any instrumental response function (IRF) by modifying Eq. 4 as appropriate; and any background photon arrival statistics or detector dark counts by modifying Eq. 5 especially relevant to in *vivo* imaging. In the Supplementary Information, Fig. S3, we tested the robustness of our method by varying the number of background photons in our data set. More significant extensions of our work, albeit generalizations that would leverage the framework at hand, would be to consider lifetime changes, due to chemical modifications of our species, over the timescale of data acquisition as may be expected in complex *in vivo* environments. ^99,100^ Another is to extend our work to analyze fluorescence lifetimes over multiple spatial locations, the purview of fluorescence lifetime imaging (FLIM) analysis.^72,101–104^ Finally, we could also generalize our proposed method to accommodate non-exponential lifetime decays if such decay probabilities are warranted by the data by modifying Eq. 5.

These, and further generalizations that can be implemented within a BNP framework, highlight the flexibility afforded by BNPs and the nature of what can be teased out from challenging data sets. Indeed, BNPs themselves suggest productive paths forward to tentatively formulate inverse strategies for challenging data sets not otherwise amenable to traditional, parametric, Bayesian analysis. ^105^

## METHOD

Here, we describe the mathematical formulation of our analysis method of time-resolved pulsed excitation single photon arrival data. For clarity we focus on measurements obtained on a fluorescence setup that use a train of identical excitation pulses. Following each pulse, one of more molecules located near the illuminated region may be excited from their ground state. As the excited molecules decay back to their ground state they may emit photons and we record the detection time. Below we describe how we analyze such recorded times.

We start from single photon detection times which consist of the raw output in a time-resolved pulsed excitation single photon arrival experiment. Similarly, these are measured based on the time difference between excitation pulses, which are time stamped, and the detection time of the first photon arriving after each pulse. ^18,39,106^ Precisely, our raw input is **Δ*t*** = (Δ*t*_1_, Δ*t*_2_, …, Δ*t_K_*) where Δ*t_k_* is the time interval between the preceding pulse’s time and the photon detection time of the *k^th^* detection. In the literature, each Δ*t_k_* is often termed micro-time. As, some pulses may not lead to a photon detection, in general the micro-times in **Δ*t*** are fewer than the total number of pulses applied during an experiment.

### Model description

We assume that, once excited, each molecule remains excited for a time period that is considerably lower (typically few nanoseconds) as compared to the time between two successive pulses (typically more than four times of the longest decay time in the sample^18^). This condition allows us to consider that any photon which is detected stems from an excitation caused by the very previous pulse and not from earlier pulses. Also, as excitation pulses in time-resolved pulsed excitation single photon arrival experiments are weak,^38,98^ and typically one in ≈ 100 pulses results in a photon detection, ^18^ we ignore, to a very good approximation, multiple photon arrivals. As the number of detected photons coming from the background is considerably lower than the number of detected photons coming from the excited molecules, typically one to ≈ 1000, we also ignore background photons. However, background photons can be dealt with straightforwardly by modifying Eq. 7 to incorporate the effect of background in the model.”

To analyze the recordings **Δ*t***, we assume that the sample contains in total *M* different molecular species that are characterized by different lifetimes *τ*_1_,…, *τ_M_*. Since molecules of each species may be excited by the pulses with different probabilities (because of different fraction of molecules contributing photons from different species), we consider a probability vector 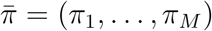 that gathers the probabilities of each species giving rise to a photon detection. Allowing *s_k_* to be a tag attaining integer values 1, …, *M*, that indicates which species triggered the *k^th^* detection, we may write

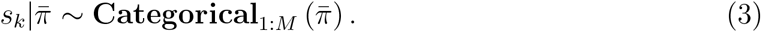

The above equation reads as follows “the tag *s_k_* given 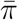 is a random variable sampled from a Categorical distribution.” The Categorical distribution is the generalization of the Bernoulli distribution which allows for more than two outcomes.^107–109^ With this convention, the lifetime of the molecule triggering the *k^th^* detection is *τ_s_k__*. Of course, the number of molecular species *M* and the precise values of the lifetimes *τ*_1_,…, *τ_M_* are unknown and our main task is to estimate them using the recordings in **Δ*t***.

For clarity, we denote with *t_pul,k_* the application time of the pulse that triggers the *k^th^* photon detection. More precisely, *t_pul,k_* is the time of the pulse’s peak. Because, in general pulses last for some non-zero duration, and so they may excite the molecules at slightly different times, we denote with *t_ext,k_* the absorption time of the molecule triggering the *k^th^* detection. Further, we denote with *t_ems,k_* the emission time of the photon triggering the *k^th^* detection. Finally, due to the measuring electronics, the detection time, which we denote with *t_det,k_*, might be different from *t_ems,k_*; see Fig. 10 for more details.

**Figure 10:**
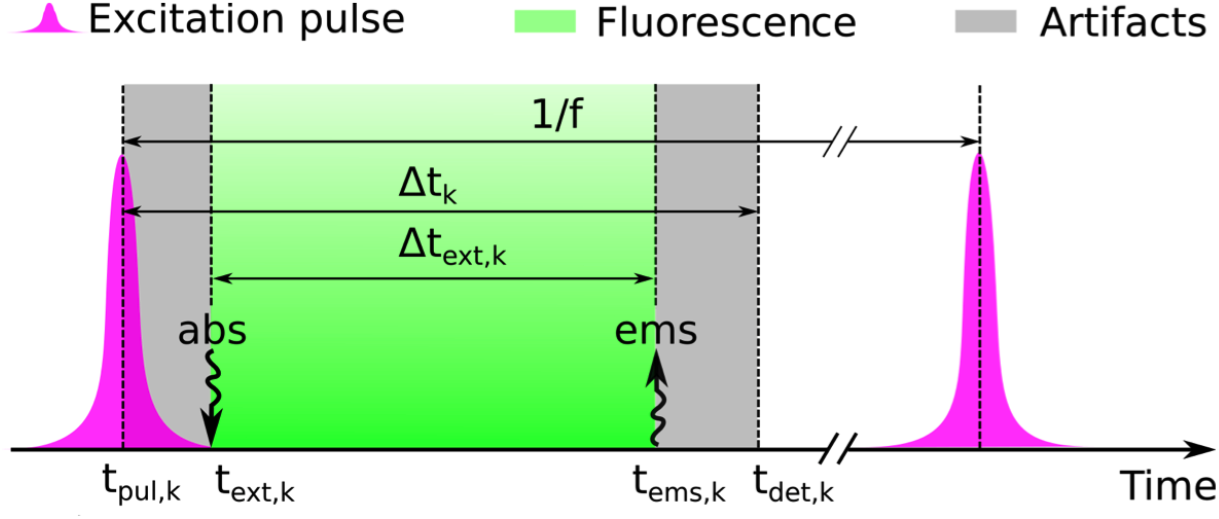
Cartoon of the factors that contribute to the recorded photon arrival times. Here, *t_pul,k_* is the time of the pulse’s peak. Since pulses last for some time, they may excite the molecules at slightly different times. As such, we denote with *t_ext,k_* the absorption time of the molecule triggering the *k^th^* detection. Moreover, we denote with *t_ems,k_* the emission time of the photon triggering the *k^th^* detection. At last, on account of electronics limitations, the detection time, which we denote with *t_det,k_*, might be different from *t_ems,k_*. Here, the artifacts shown in gray originate from two sources: the left gray-shaded region is due to the width of the pulse which leads to variation in the time of the molecular excitation, and the right gray-shaded region arises from the camera-dependent detection uncertainty. The time during which the fluorophore is excited (fluorescence lifetime) is shown in green.

With this convention, our measured output consists of the time lags Δ*t_k_* = *t_det,k_* – *t_pul,k_*. These time lags include: (i) the time until absorption occurs, *t_ext,k_* – *t_pul,k_*; (ii) the time until fluorescence emission occurs, *t_ems,k_* – *t_ext,k_*; (iii) delays and errors introduced by the measuring electronic devices, *t_det,k_* – *t_ems,k_*. Below, we denote the middle period with Δ*t_ext,k_* = *t_ems,k_* – *t_ext,k_*; while, we denote with Δ*t_err,k_* – (*t_ext,k_* – *t_pul,k_*) + (*t_det,k_* – *t_ems,k_*) the sum of the others. From these two, Δ*t_ext,k_* is the time the molecule spends in the excited state; while, Δ*t_err,k_* gathers any artifacts caused by our setup either in the excitation or detection pathway. The advantages of considering these two periods separately, as we explain below, is that (i) these represent independent physical processes, and (ii) each one is theoretically and experimentally characterized well. ^18^

In particular, Δ*t_err,k_* is characterized by the instrument response function (IRF) that, in each set-up, is readily obtained with calibration measurements.^18^ In this study, we approximate the IRF as a Gaussian

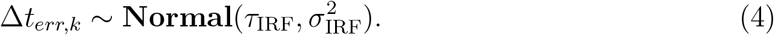

In this approximation, *τ*_IRF_ is the IRF’s peak time and *σ*_IRF_ = FWHM/2.355 where FWHM is the IRF’s full-width-at-half-maximum. In the Supplementary Information, we explain the IRF’s calibration in detail.

Upon excitation, the time the molecule remains excited, Δ*t_ext,k_*, is memoryless,^18^ and so it follows the exponential distribution. Therefore,

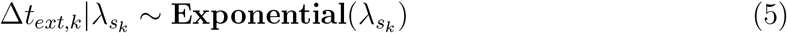

where λ*_s_k__* is the inverse lifetime of the molecule triggering the detection of Δ*t_ext,k_*. Of course, the inverse lifetime depends upon the lifetime by λ*_s_k__* = 1/*τ_s_k__*.

Because Δ*t_ext,k_* and Δ*t_err,k_* are independent variables, the statistics of our measurements, which are given by Δ*t_k_* = Δ*t_ext,k_* + Δ*t_err,k_*, follow

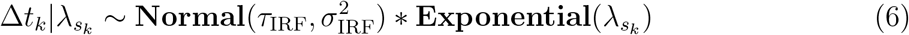

where ∗ denotes a convolution,^110^ and specifically has the probability density

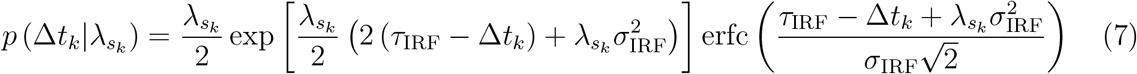

where erfc(·) denotes the complementary error function. In the Supplementary Information, we show analytically how Eq. (7) arises from Eqs. (4) and (5).

In the next section we describe how Eqs. (3) and (7) can be used in conjunction with BNP to obtain the estimates we are after.

### Model inference

All quantities which we wish to infer, for example the species inverse lifetimes λ_1_,…, λ_*M*_ and excitation probabilities in 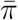, are represented by model variables in the preceding formulation. We infer values for these variables within the Bayesian paradigm. ^80,82,84^ Accordingly, on the inverse lifetimes we place independent priors

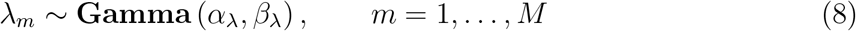

that ensure strictly positive values. Here, for convenience only, we consider priors on inverse lifetimes where 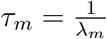 is the molecular lifetime and λ*_m_* is the inverse lifetime of species *m*. As the total number of species contributing photon detections in an experiment is unknown, we consider a symmetric Dirichlet prior^80,83^ (which is conjugate to the Categorical) on 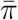 of the form

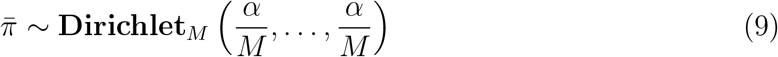

where *α* is a positive scalar hyper-parameter. A graphical summary of the whole formulation is shown on Fig. 11.

**Figure 11:**
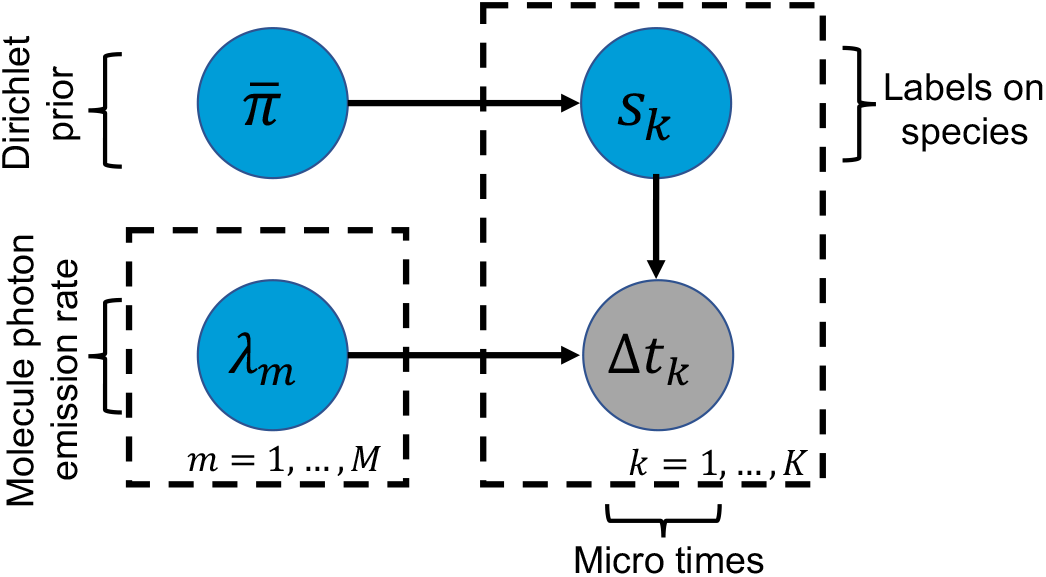
Graphical representation of the proposed model. A simple graphical representation of the model, where Δ*t_k_* is the micro time *k* with *k* = 1,…, *K*. The inverse lifetime of species m is shown by λ_*m*_, *m* = 1,…, *M*. The label *s_k_* tells us which of the species is contributing the *k^th^* photon. In the graphical model, the measured data are denoted by grey shaded circles and the model variables, which require priors, are designated by blue circles. Each one of the labels has a prior which is a Dirichlet probability 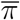.

The distribution in Eq. (9) ensures that 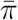 are valid probability vectors. Further, Eq. (9) is specifically chosen to allow for a large, *M* → ∞, number of species. This is particularly important because the total number of molecular species contributing to the detections in TCSPC or FLIM experiments are typically unknown, and so choosing a finite *M* may lead to under-fitting. Specifically, at the limiting case *M* → ∞, the prior on Eq. (9), combined with Eq. (3), results in a Dirichlet process.^75,83,111,112^ In other words, provided *M* is sufficiently large, the estimates obtained through our model are independent of the particular value chosen (*i.e*., overfitting cannot occur).

With the nonparametric model just presented, although the total number of model molecular species is infinite, the actual number of molecular species contributing photons to the measurements is finite. Specifically, the number of contributing species coincides with the number of different tags *s_k_* associated with **Δ*t***. In other words, instead of asking *how many species contribute to the measurements?*, with our model, we ask *how many of the represented species actually contribute at least one photon?* Further, instead of asking *what are the lifetimes of these species?* we ask *what are the lifetimes of the species contributing at least one photon?* Of course, as we estimate inverse lifetimes instead of lifetimes, we obtain the latter by *τ_m_* = 1/λ_*m*_.

With these priors, we form 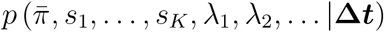 which is the joint posterior probability distribution that includes all unknown variables. To compute this posterior, we develop a Markov Chain Monte Carlo (MCMC) scheme^84,113^ that generates pseudo-random samples with the appropriate statistics. The scheme is described in the Supplementary Information and a working implementation is also provided.

### Acquisition of Synthetic Data

The synthetic data presented in this study are obtained by standard pseudo-random computer simulations^114–118^ that simulate a common fluorescence lifetime imaging modality with a conventional single-spot confocal setup. Further, in the simulations we consider confocal regions created with pulsed excitation. To generate data mimicking as closely as possible the measurements obtained in real experiments, we simulate freely diffusing molecules of different species characterized by different diffusion coefficients and lifetimes. Details and parameter choices are provide in the Supplementary Information, Tables S2 and S3.

### Acquisition of Experiment Data

The synthetic data presented in this study are obtained as described below.

#### Sample preparation

Sample solutions of Rhodamine B (Rhod-B, Wako Pure Chemical Industries), Rhodamine 6G (Rhod-6G, Sigma-Aldrich), and tetramethylrhodamine-5-maleimide (TMR, Invitrogen), and Cy3 monofunctional NHS-ester (Cy3, GE Healthcare) were prepared with Milli-Q water at 1 *μ*M concentration. Nonionic surfactant (0.01% Triton X-100) and 2 mM Trolox were added to prevent adsorption of dye molecules to the glass surface and reduce photophysical artifacts, respectively.

#### Experiments

Fluorescence lifetime measurements were carried out using a confocal fluorescence microscope with super continuum laser (Fianium SC-400-4, frequency of 40 MHz). The output of the laser was filtered by a bandpass filter (Chroma Technology D525/30 m), and focused onto the sample solution using a 60× objective lens (Nikon Plan Apo IR) with NA of 1.27. The excitation power was set to be 0.3 *μ*W at the entrance port of the microscope. Fluorescence photons ware collected by the same objective lens and guided through a confocal pinhole as well as a bandpass filter (Chroma Technology D585/40 m), and then detected by a hybrid detector (Becker & Hickl HPM-100-40-C). For each photon signal detected, the routing information was appended by a router (Becker & Hickl HRT-82). The arrival time of the photon was measured by a TCSPC module (Becker & Hickl SPC-140) with the time-tagging mode. ^37^ The time resolution was evaluated by detecting the scattering of the incident laser light at a cover glass, and it was typically 180 ps at full width half maximum.

## ACKNOWLEDGEMENTS

S.P. acknowledges support from the NIH NIGMS; R01GM134426 “Theoretical models of single molecule dynamics from minimal photon numbers” for the contributions and support of S.J. and R01GM130745 “A Bayesian nonparametric approach to superresolved tracking of multiple molecules” for the contributions and support from I.S. T.T. acknowledges support from JSPS KAKENHI Grant Number JP19F19340.

## AUTHOR CONTRIBUTIONS

MT and SJ analyzed data and developed analysis software; MT, SJ, IS developed computational tools; WH, KI, TT contributed experimental data; MT, SJ, IS, SP conceived research; SP oversaw all aspects of the projects.

## SUPPLEMENTARY INFORMATION

In this supplement, we present additional analyses and technical details expanding upon the material presented in the main text. These include: (i) additional analysis of synthetic and experimental traces that include the estimation of lifetimes and the fraction of different species contributing photons; (ii) additional details on the theoretical approaches used; and (iii) a complete description of the inference framework developed that includes choices for prior probability distributions and a computational implementation. Moreover, in our BNP analysis, we do not pre-specify the number of species, we learn them.

### Additional results

#### Analysis of additional synthetic data

In the main text we focused on the estimation of: lifetime, *τ*, with values less than 10 ns which are typical lifetime values in *in vivo* applications.^99^ Here, we explore broader parameter ranges from freely diffusive molecules, Figs. S1 and S2 to the case when we have different background photons, Fig. S3, which we evaluated our method respect to different background levels to see how it behaves with different number background photons. Moreover, we evaluated our method for cases with more than two species, Figs. S4 and S5, and estimate the fraction of molecules contributing photons from different species, Fig. S6, that we explain in the main text in Section “Mixtures of different species contributing photons”.

**Figure S1:**
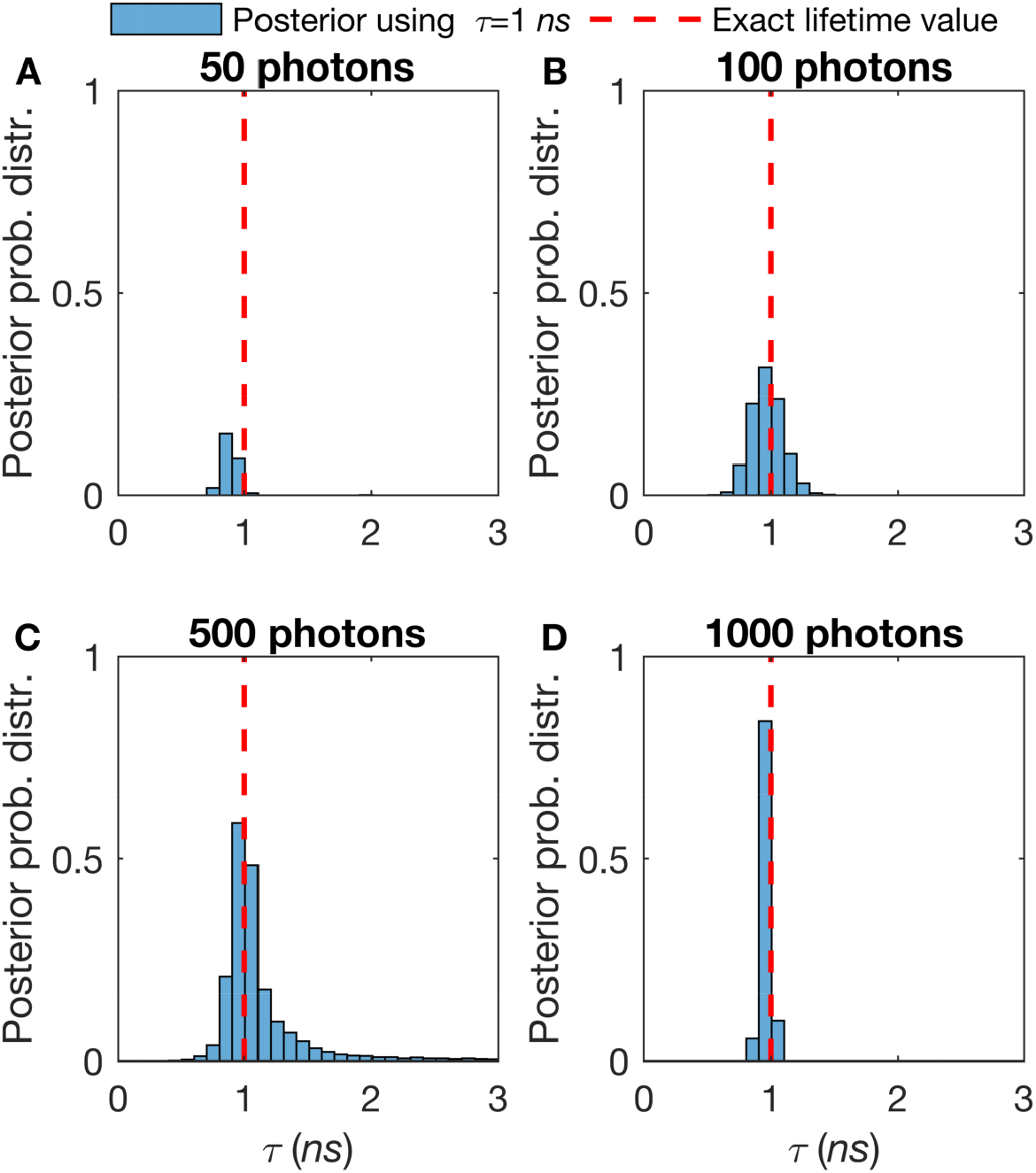
Effect of the number of detected photons on a single diffusive molecular lifetime estimate. The more photons per unit time, the sharper the lifetime estimate. Here, we work on single species lifetime while all molecules are diffusing with diffusion coefficient, *D* = 10 *μm*^2^/*s*. The synthetic trace is generated using *τ* =1 *ns*. We start with 50 photons (A) and gradually increase the number of photons that we incorporate into the analysis to 100 (B), 500 (C), and 1000 (D) photons. The excitation pulses occur at a frequency of 40 *MHz* and we assume that these pulses assume a Gaussian shape with standard deviation of 0.1 *ns*. The ground truth for the lifetimes are known (as this is synthetic data) and they are shown by red-dashed lines.

**Figure S2:**
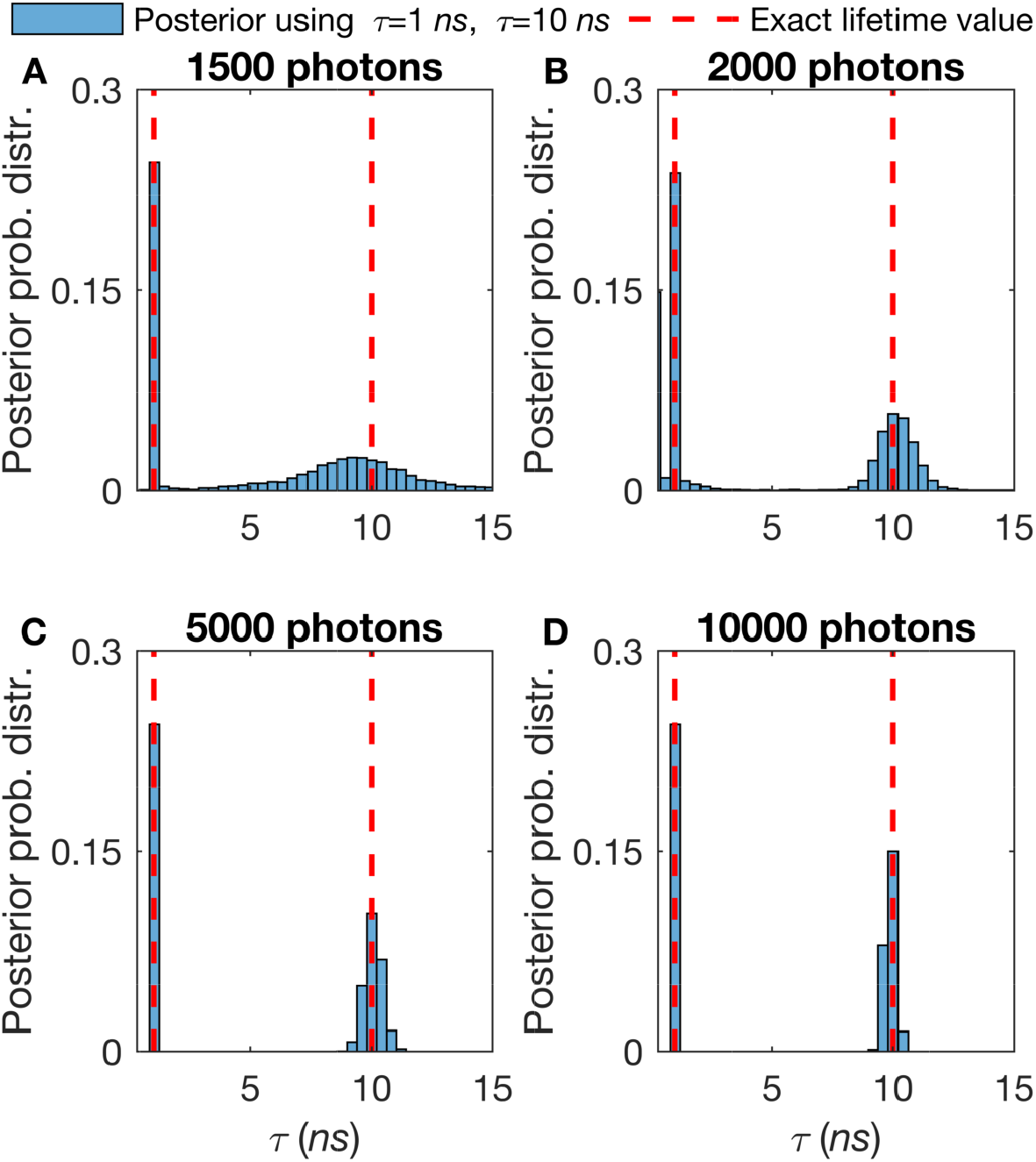
Effect of the number of detected photons on a double diffusive molecular lifetime estimation. The more photons per unit time and thus the sharper estimation of lifetime. Here, we work on single species lifetime while all molecules are diffusing with diffusion coefficient, *D* =10 *μm*^2^/*s*. The synthetic trace generated by *τ* =1 *ns* and *τ* =10 *ns*. We start with 1500 photons (A) and gradually increase the number of photons that we incorporate into the analysis to 2000 (B), 5000 (C), and 10000 (D) photons. Here, all other features such as the frequency of acquisition and width of pulse are the same as in Fig. S1. Also, we follow the same red-dashed line convention as in 3.

**Figure S3:**
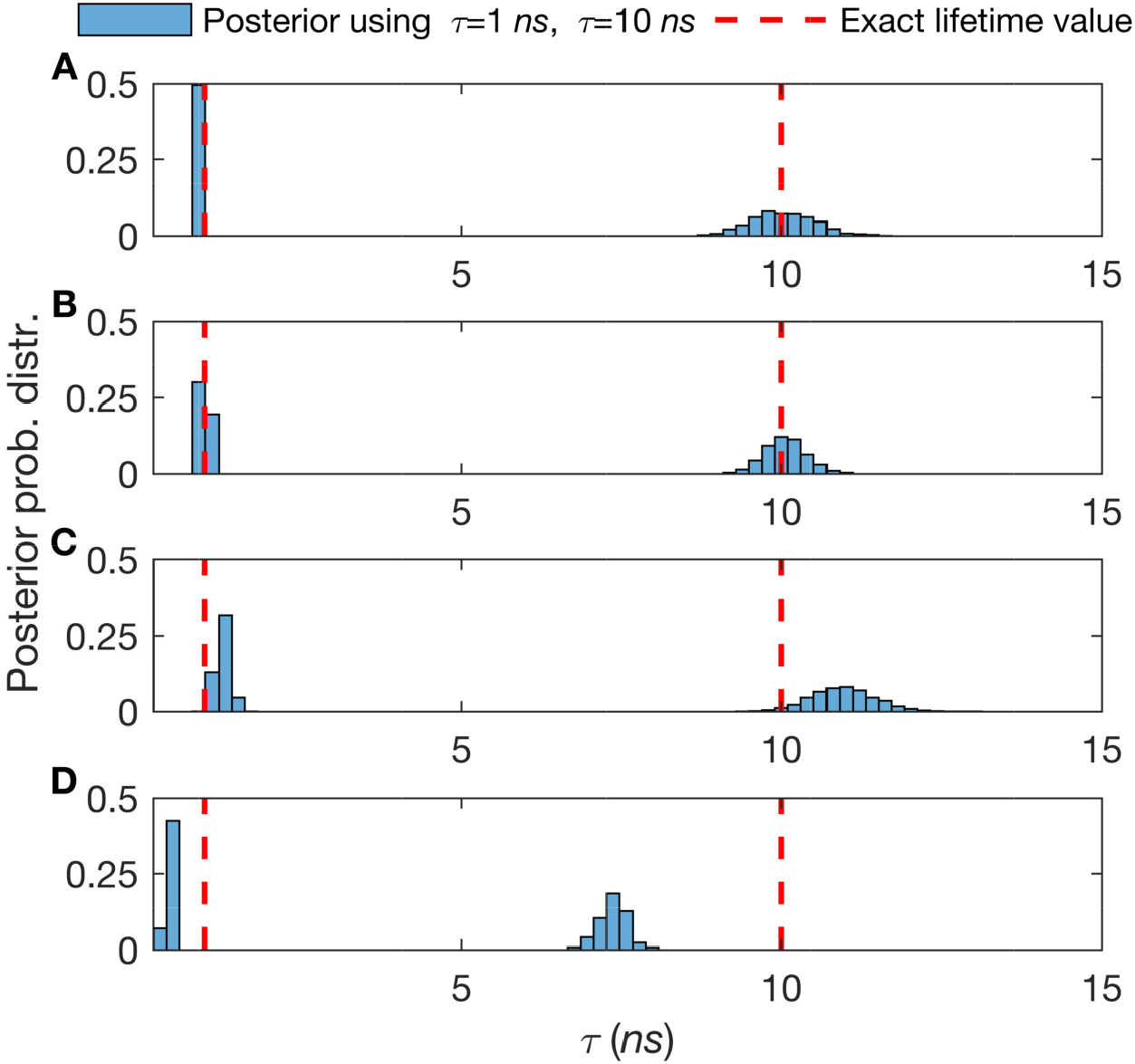
Effect of the number of background photons on a double diffusive molecular lifetimes estimation. The more background photons per unit time, the poorer the lifetime estimate. Here, we work on double species lifetime while all molecules are diffusing with diffusion coefficient, *D* = 10 *μm*^2^/*s*. The synthetic trace generated by *τ* = 1 *ns* and *τ* = 10 *ns* with total 3000 photons. We start with 3 background photons (A) and gradually increase the number of photons that we incorporate into the analysis to 30 (B), 150 (C), and 300 (D) photons. Here, all other features such as the frequency of acquisition and width of pulse are the same as in Fig. S1. Also, we follow the same red-dashed line convention as in 3.

**Figure S4:**
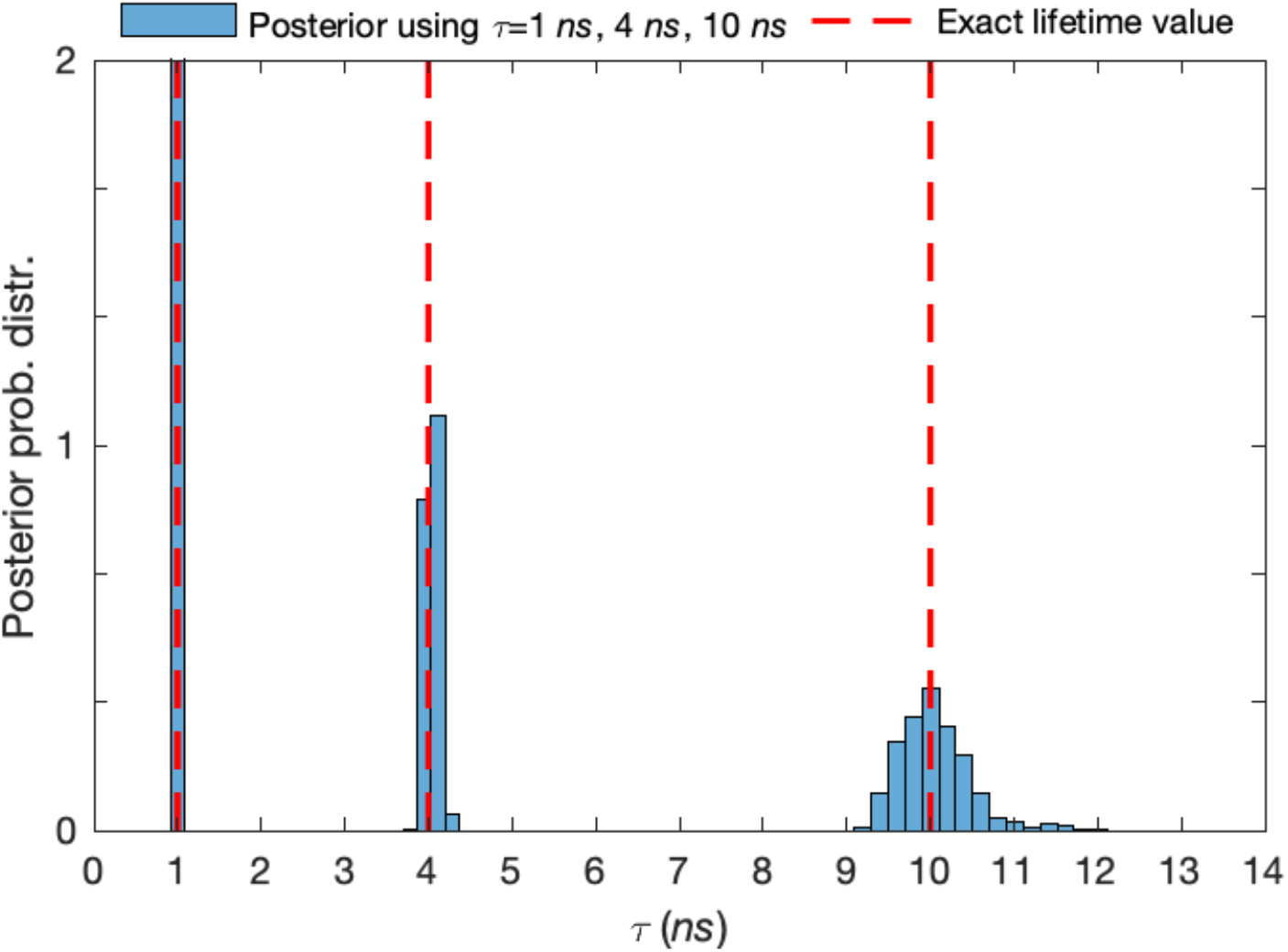
Lifetime estimates with three different species using synthetic data. Here, we generate a synthetic trace with three species having lifetimes *τ* =1 *ns*, *τ* = 4 *ns* and *τ* =10 *ns* with equal fraction of molecules contributing photons from different species (33% for each of them) and analyze a total of 2 × 10^5^ photon arrivals. Here, all other features such as the frequency of acquisition and width of pulse are the same as in Fig. S1. Also, we follow the same red-dashed line convention as in 3.

**Figure S5:**
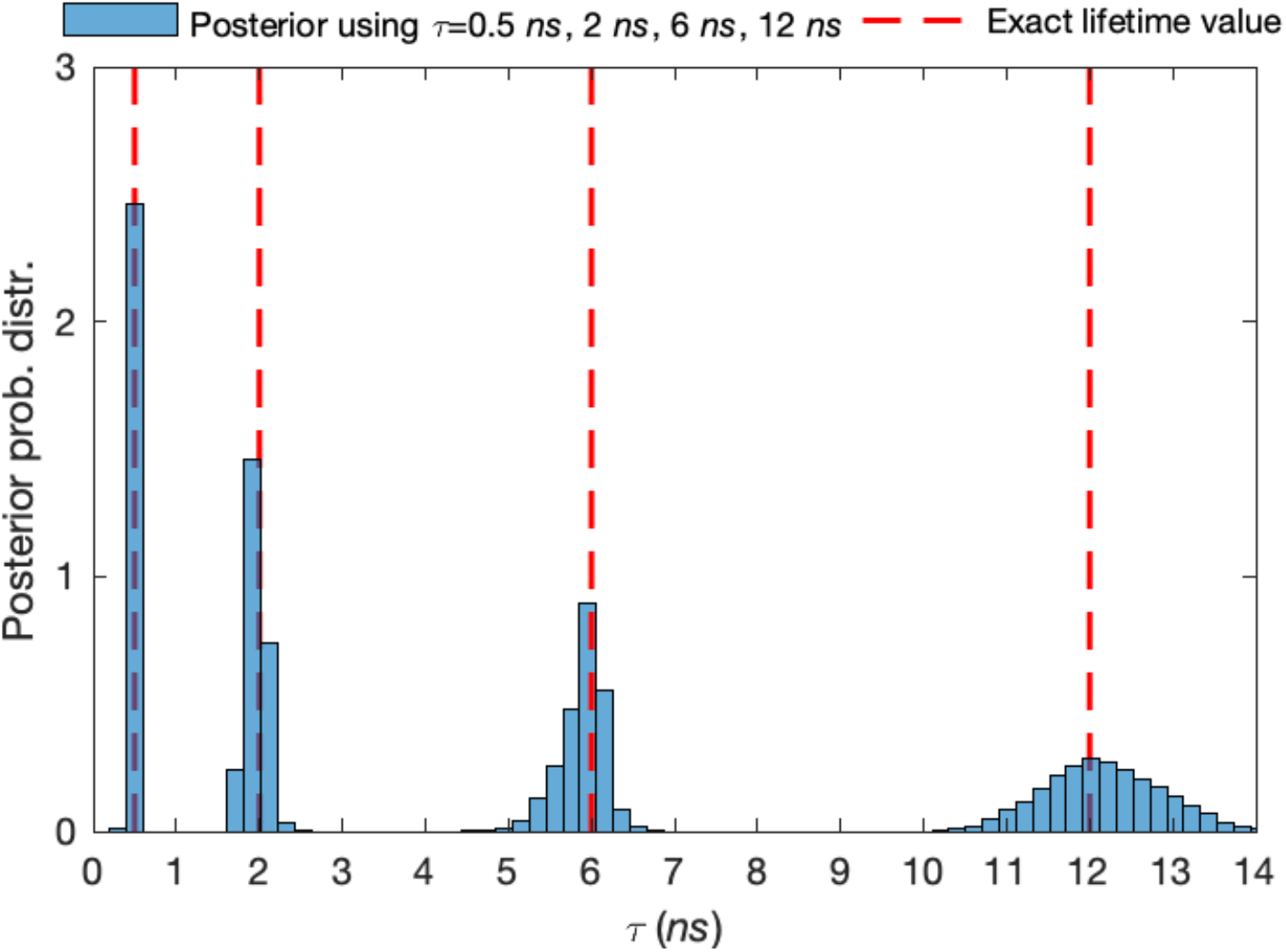
Lifetime estimates with four different species in synthetic data. Here, we work with four species lifetimes while all molecules are immobilized. The synthetic trace generated by *τ* = 0.5 *ns*, *τ* = 2 *ns*, *τ* = 6 *ns* and *τ* =12 *ns* with equal fraction of molecules of each species (*i.e*., set at 25%) for each of them and analyze a total of 3 × 10^5^ photon arrivals. Here, all other features such as the frequency of acquisition and width of pulse are the same as in Fig. S1. Also, we follow the same red-dashed line convention as in 3.

**Figure S6:**
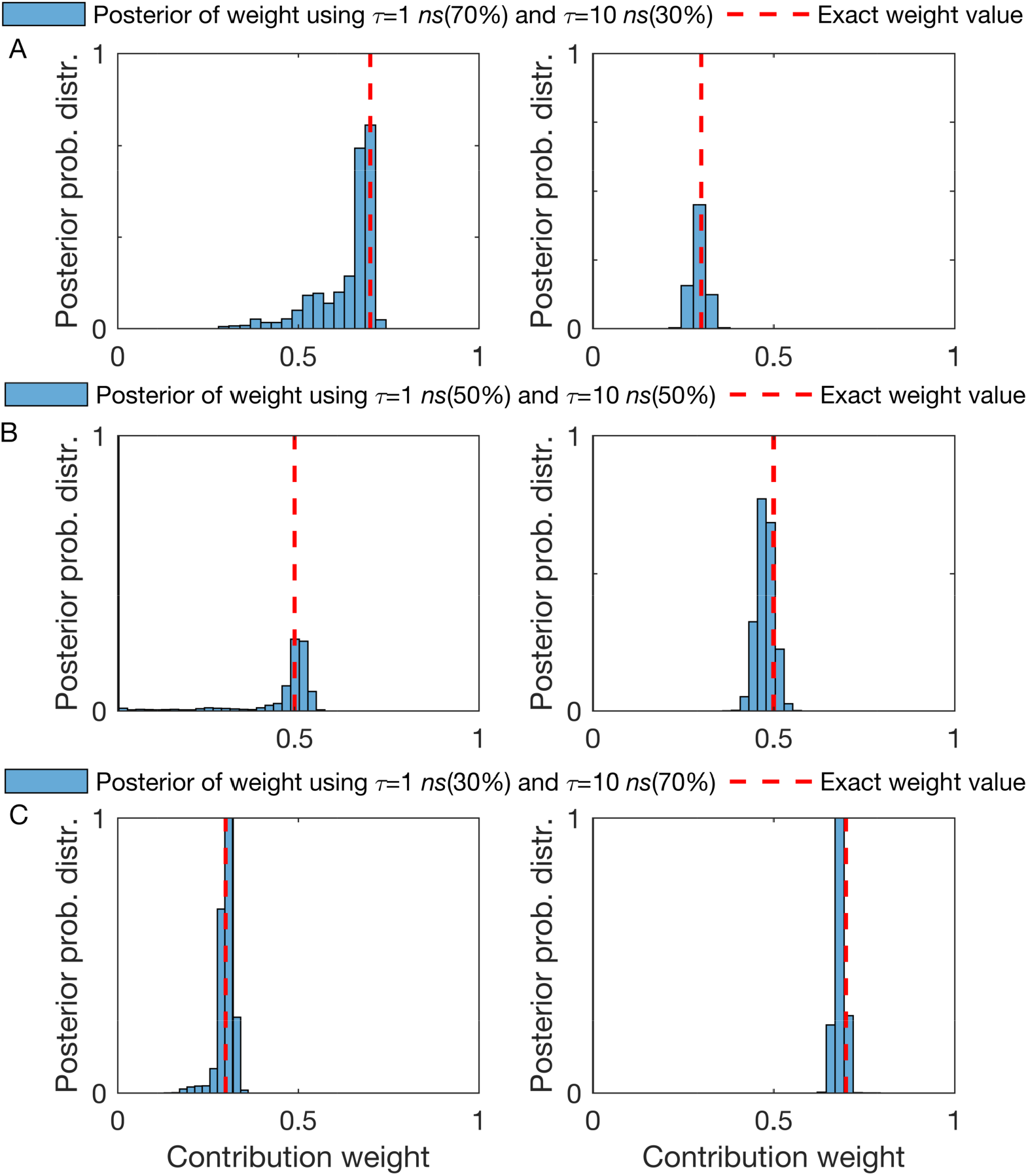
Estimation of the fraction of molecules contributing photons from different species. (A-C) Using the same synthetic traces as in Fig. 5, the posterior probability distribution over the fraction of molecules contributing photons from different species (weight) with lifetimes of 1 *ns* and 10 *ns*, 3000 total number of detected photons analyzed and fractions of chemical species of 70% – 30%, 50% – 50% and 30% – 70% respectively. Here, all other features such as the frequency of acquisition and width of pulse are the same as in Fig. S1. Also, we follow the same red-dashed line convention as in 3.

#### Analysis of additional experimental data

Here, we used real measurements, obtained as explained in the method section, from different fluorescent dyes, namely Cy3, TMR, Rhod-B, and Rhod-6G. In Fig. S7 we considered a mixture of all four species. In Fig. S8 we show that we can correctly identify the fraction of molecules contributing photons from different species.

**Figure S7:**
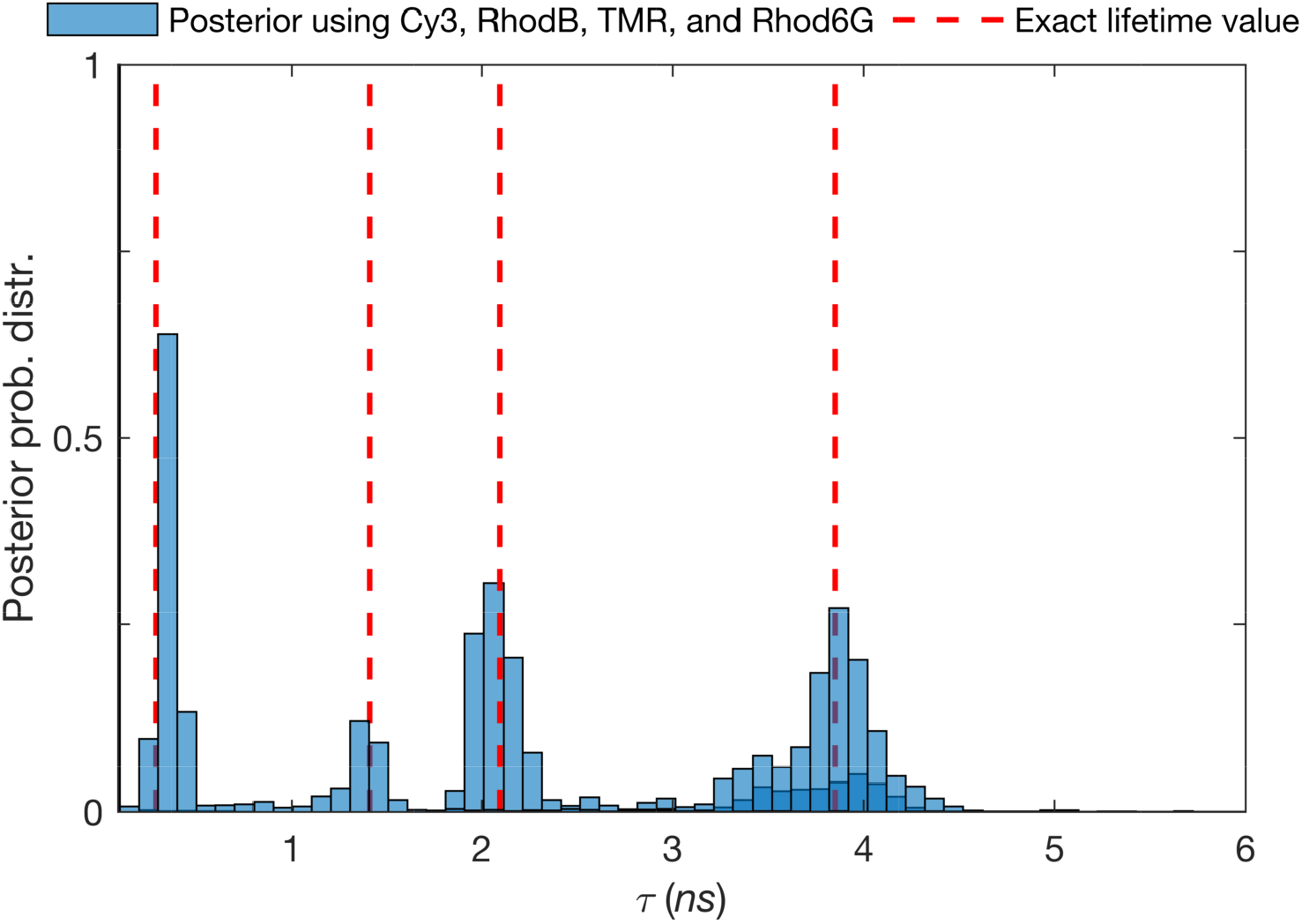
Lifetime estimates for the case of four different species from experimental data. Here, we work on four species lifetimes while all molecules are immobilized. The experimental trace generated by four different dyes including Cy3, Rhod-B, TMR, and Rhod-6G with a total of ≈ 3 × 10^5^ photon arrivals analyzed. The excitation pulses occur with a frequency of 40 *MHz* and we assume that these pulses are modeled by a Gaussian with a standard deviation of 0.1 *ns*. The ground truth estimates for the lifetimes are determined using the whole trace which includes total 1.4 × 10^6^ photon arrivals and they are shown by red-dashed lines.

**Figure S8:**
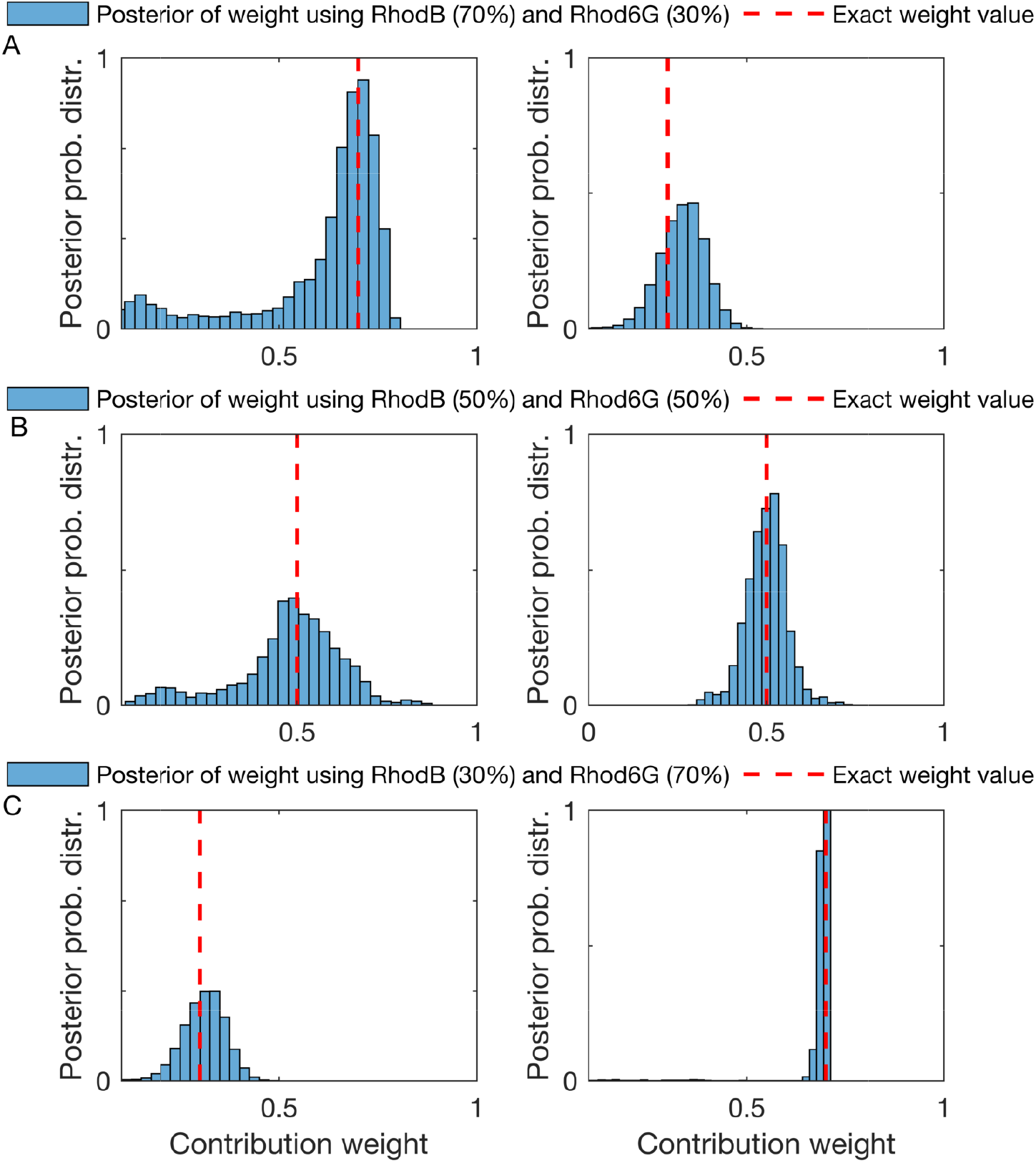
Estimation of the different fraction of molecules contributing photons from different species from experimental data. (A-C) Using the same traces as Fig. 9, the posterior probability distributions for the fraction of chemical species contributing photons for experimental dyes, RhodB and Rhod6G, with a total of ≈ 3000 total number of detected photons and fraction of chemical species contributing photons of 70% – 30%, 50% – 50% and 30% – 70% respectively. The excitation pulses happen at a frequency of 40 *MHz* and we consider them to have a Gaussian shape with a standard deviation of 0.1 *ns*. What we treat as ground truth lifetime estimates (as we do not have real ground truths for experimental data) are determined using the whole trace which includes a total of 1.4 × 10^6^ photon arrivals and they are shown by red-dashed lines.

### Brief description of phasor plots analysis

#### Time domain

In typical time-domain lifetime imaging, a pulsed laser is used to excite the sample periodically, causing fluorescence emission for those pulses where a molecule is excited and decays back to the ground state radiatively. Experimentally, based on the data we presented, this is typically 1 in 40 pulses.^18^

From Eq. 1, fluorescence species with M different lifetimes have exponentially decaying intensities

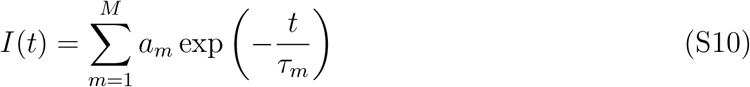

with fluorescence lifetimes *τ_m_* and weights, *a_m_*. In an ideal scenario, a fluorophore is excited with an exceedingly thin (Dirac-shaped) laser pulse at time *t* = 0. Its initial intensity is therefore *I*(*t* < 0) = 0. As excitation pulses are not infinitely sharp and detectors exhibit delays, the recorded signal, 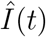, is the convolution of its fluorescence intensity *I*(*t*) with the instrumental response function (IRF);^119,120^ see Eq. S21.

#### Frequency Domain

Frequency-domain experiments constitute an alternative way to measure excited state life-times. In this case, the sample is excited with an intensity-modulated light, typically a sine-wave.^18^ When a fluorescent sample is excited in this way, the emission intensity follows a shifted modulation (*m*) pattern with the phase shift (*ϕ*) and peak height that both encode information on the excited state lifetime.^18^

**Figure S9:**
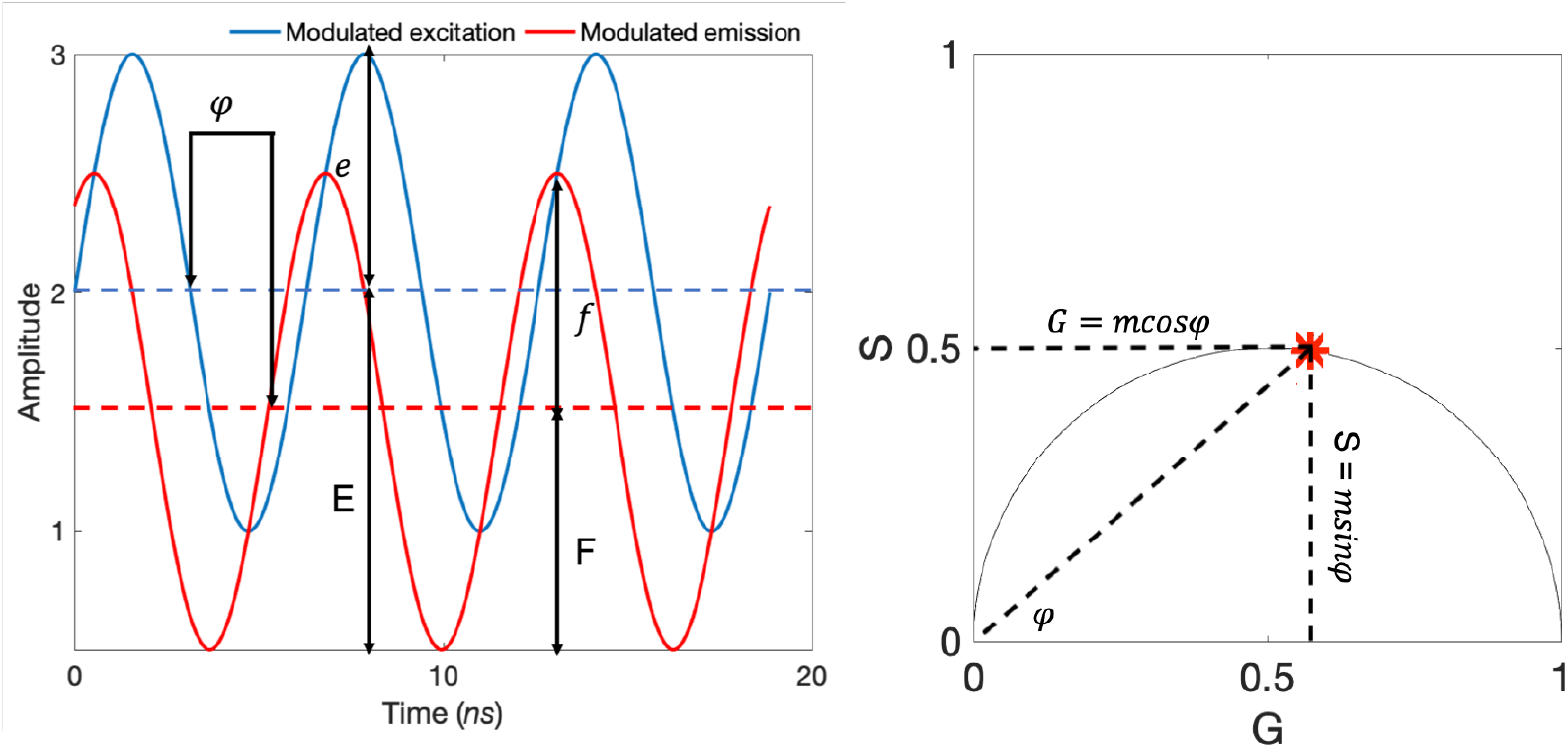
Frequency-domain analysis. Mapping frequency-domain modulated emission (left) into a phasor plot representation (right).

The modulation of the excitation is given by 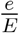, where *e* is the average intensity and *E* is the peak-to-peak height of the incident light (Fig. S9). The modulation of the emission is defined similarly, 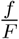, except using the intensities of the emission (Fig. S9). The shifted modulation between emission and excitation, *m* = (*f/F*)/(*e/E*). The other experimental observable is the phase shift, (*ϕ*) which is the phase difference between excitation and emission. Both phase shift (*ϕ*) and the shifted modulation between emission and excitation (*m*) can be employed to calculate the lifetime using

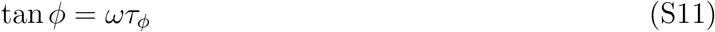

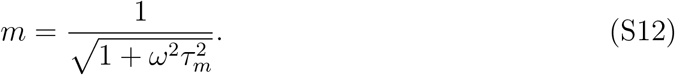

These expressions can be also be used to calculate the phase (*τ_ϕ_*) and shifted modulation (*τ_m_*) lifetimes for the curves shown in Fig. S9. If the intensity decay is a single exponential, then Eqs. S11 and S12 yield the correct lifetime. In this case, both *τ_ϕ_* and *τ_m_* are equal. For more than one species, these two are not the same and details are discussed in Ref.^18^

Along these same lines, lifetimes can also be determined using a phasor approach first introduced by Jameson et al.^121^

Briefly, we introduce the pair of conjugate variables *G* and *S* (termed phase coordinates) where

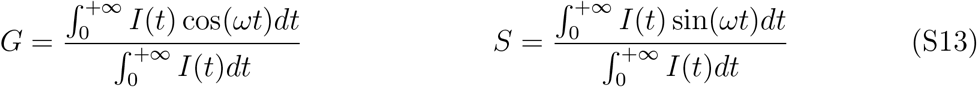

and where *I*(*t*) is the photon intensity.^68,119^ In the case of single exponential 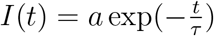, the coordinates of the phasor are given by

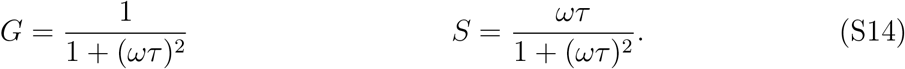

### IRF approximation

To incorporate the effect of the IRF in our analysis, we approximate the IRF with a Gaussian function;^31^ see Fig. S10. Centrally symmetric pulses such as the Gaussian, are obtained from electronics as used in most modern instruments.^33^ However, non-symmetrical IRFs could be handled by proper modifications to Eq. 4 in the main text.

**Figure S10:**
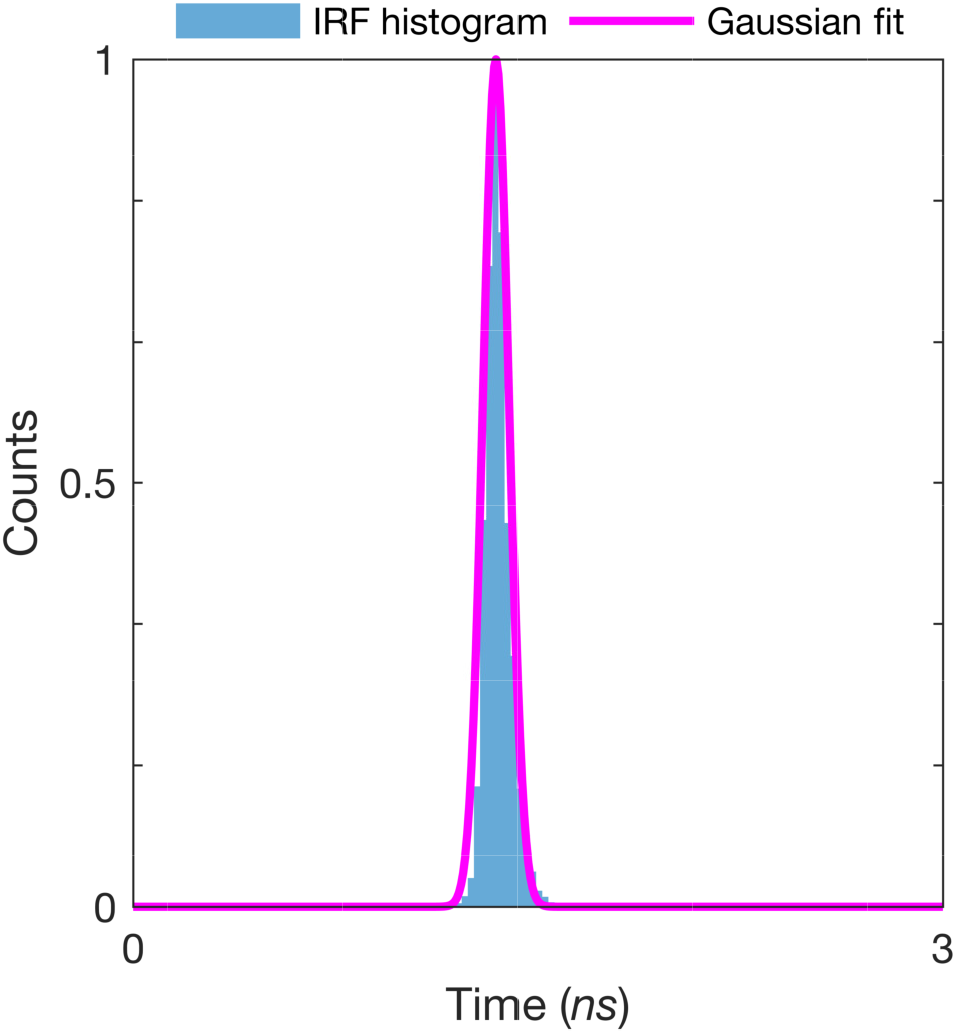
The actual IRF (blue color) fitted with a Gaussian function (magenta color). The fitted IRF is used for the analysis of all experimental data.

### Description of the pulsed excitation and microtimes simulation

To simulate experimentally realistic microtimes, for mobile particles, we simulate diffusive molecules which freely traverse through an illuminated confocal volume. We define periodic boundaries (±*L_x_*, ±*L_y_*, ±*L_z_*) which are much larger than the confocal radii to maintain a constant concentration of molecules. The confocal volume itself is pulsed on and off and the probability of excitation of a molecule depends on its location within that volume during the pulse. Here we consider the confocal volume (the combined excitation and emission point spread function, PSF) to be a 3D Gaussian, with radii of *ω_x_* = 0.3 *μ*m, *ω_y_* = 0.3 *μ*m, *ω_z_* = 3.5 *μ*m and centered at the point of origin. The precise formula for this PSF is

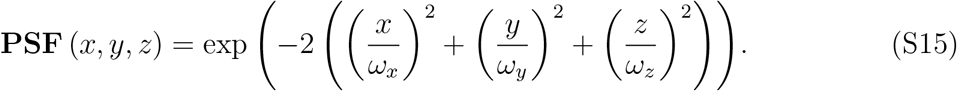

**Figure S11:**
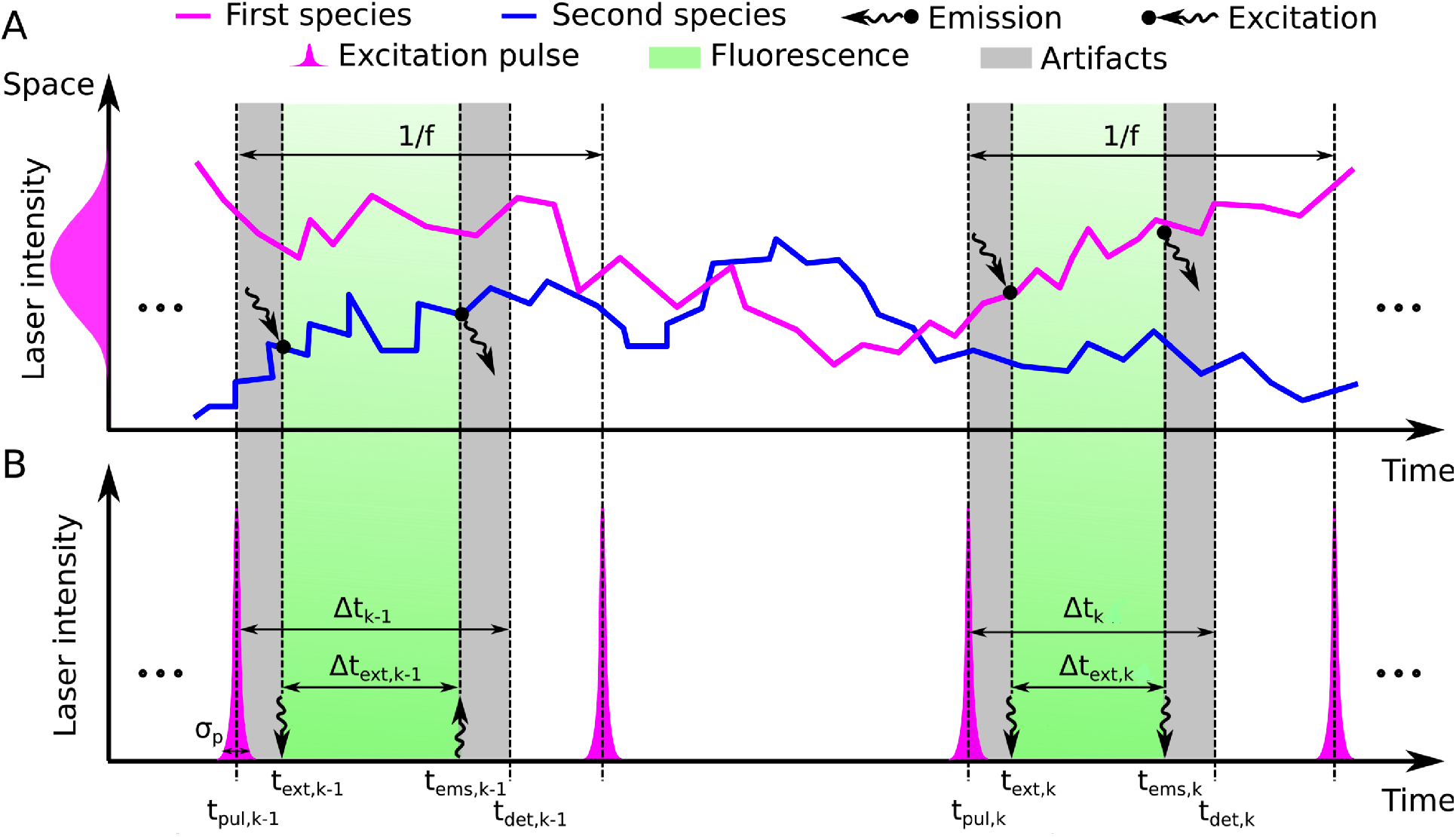
Pictorial representation of the experimental setup a sample with a mixture of two species. (A) The Brownian motion of two species in space versus time. Excitation and emission points are shown with different arrows. (B) Micro-times are the time between the peak of the pulse *t_pul,k_* that trigger the *k^th^* photon detection and detection time *t_det,k_*. The time between the excitation *t_ext,k_* and emission *t_ems,k_* of the molecule, Δ*t_ext,k_* follows the molecular lifetime. The gray and green-shaded regions are described in Fig. 10.

So, the emission that received by molecule *n* of the *m^th^* is species

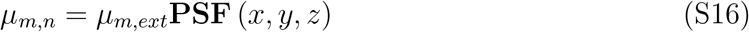

where, *μ_m,ext_* is the maximum excitation rate of the molecule *n* of species *m* which occurs when the molecule is at the center of the confocal volume.^122^

Assuming that molecules do not move significantly over the duration of the pulse (of typical width 0.1 ns^123^), the probability of excitation of molecule *n* of species *m* is *q_m,n_ = μ_m,n_δt_p_* where, *δt_p_* is the duration of the pulse. So, for any pulse excitation, we need to determine if the *n^th^* molecule of species *m* is excited or not. We define the variable *b_m,n_* to be either 1 or 0 if the molecule emits or does not emit a photon and consider this variable to be Bernoulli distributed

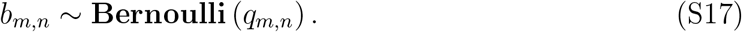

At the end, when a molecule is excited by each pulse *b_m,n_* = 1, we need to consider the delays and errors introduced by the measuring electronic devices, *t_det,k_ − t_ems,k_*. Since, we consider these errors follow a normal distribution, and the excitation time is normal distributed as well, we denote both effects with Δ*t_err,k_* = (*t_ext,k_ − t_pul,k_*) + (*t_det,k_ − t_ems,k_*) and as the result, we sample it from a normal distribution

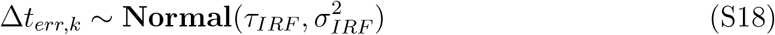

where *τ_IRF_* is the mean of IRF and *σ_IRF_* is the standard deviation of the IRF (see Eq. 4 for comparison). In this simulation we considered 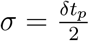 as the width of the pulse.

After sampling the error time, we sample the emission time of each molecule from the exponential distribution with corresponding inverse lifetime belongs to species *m*

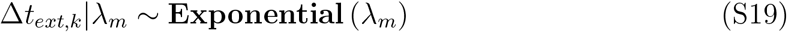

and as we have shown in the Fig. S11 the detection time of each molecule will be sum of these two times

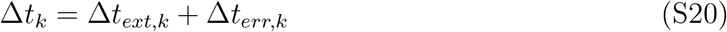

which is determined by the convolution of emission profile, Eq. S18, and excitation pulse, Eq. S19.

### Derivation of model likelihood

As we mentioned in the main text, Section “Model description”, measurements Δ*t_k_* = Δ*t_ext,k_* + Δ*t_err,k_*, follow

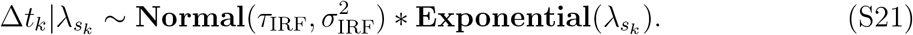

In this case we have

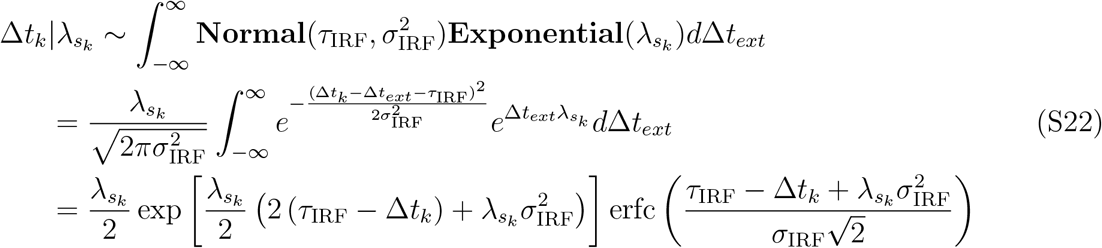

where erfc(·) denotes the complementary error function.

### Detailed description of the inference framework

#### Description of prior probability distributions

Within the Bayesian approach, all unknown model parameters need priors. The model parameters in our framework that require priors are: the inverse lifetimes {λ_*m*_}_*m*_; labels on each species 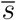; and probability on the labels of species 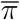 (fraction of molecules contributing photons from different species). Our choices of priors are described below.

##### Inverse lifetimes, {λ_*m*_}_*m*_

Here we are faced with different species which have different lifetimes. For convenience, we consider inverse lifetimes instead of lifetimes, 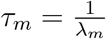, where the *τ_m_* is the molecular lifetime and λ_*m*_ is the inverse lifetime of species *m*.

To learn inverse lifetimes, and to guarantee that their sampled values in our formulation attain only positive values, we place a Gamma distribution prior over them as follows

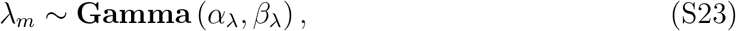

where, *α*_λ_ and *β*_λ_ are prior parameters.

##### Weights, 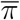

The weight on each species comes from the Dirichlet distribution

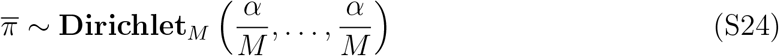

where *α* is the scalar parameter of the Dirichlet distribution.^75,124^ This prior is conjugate to the labeled species, *s_k_*, which simplifies the computations shown below. The Dirichlet distribution is an important multivariate continuous distribution in Bayesian statistics which is a multivariate generalization of the Beta distribution and, conveniently, conjugate to the Categorical.^125^

#### Labels on each species, *s_k_*

Since we have many species, we define a label for each molecule which will tell us that molecule belongs to which species

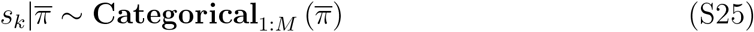

where 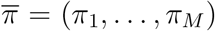 is the weight on each species. In other words, *π_m_* is the fraction of photons which species *m* contributes to the data.

### Summary of model equations

For concreteness, below we summarize all equations used in our framework, including a complete list of priors.

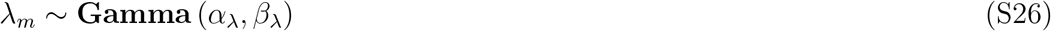

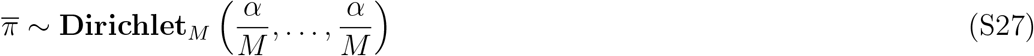

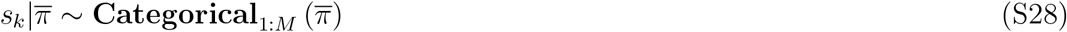

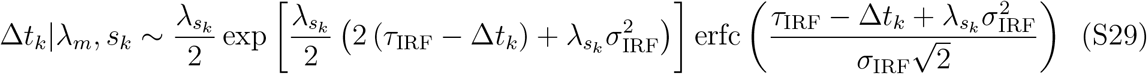

### Inverse problem

Within the Bayesian paradigm, our goal is to sample from the following posterior probability distribution 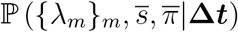. Since, it is not possible to directly compute this distribution, we will sample the random variables 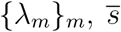, and 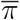 from their conditional distributions through a Gibbs sampling scheme.^80,81,84,113,126^ Accordingly, posterior samples are generated by updating each one of the variables involved sequentially by sampling conditioned on all other variables and the measurements **Δ*t***.

Conceptually, the steps involved in the generation of each posterior sample 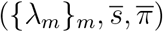 are:

Update the weights on each species 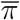
Update the labels on species 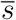
Update the inverse lifetimes {λ_*m*_}_*m*_.

#### Sampling of the weights 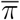

To update the weights of the labels on the species 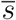, we sample them from the corresponding conditional probability 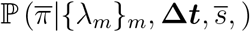, which simplifies to 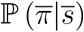.

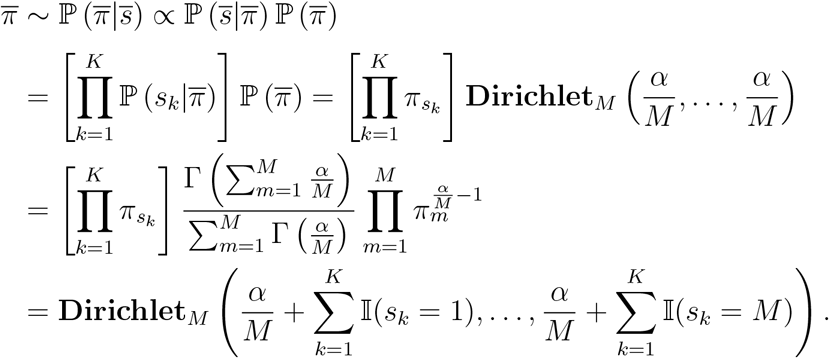

#### Sampling of the labels 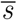

To sample the labels on species, we sample them from the conditional probability distribution 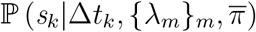 as follows

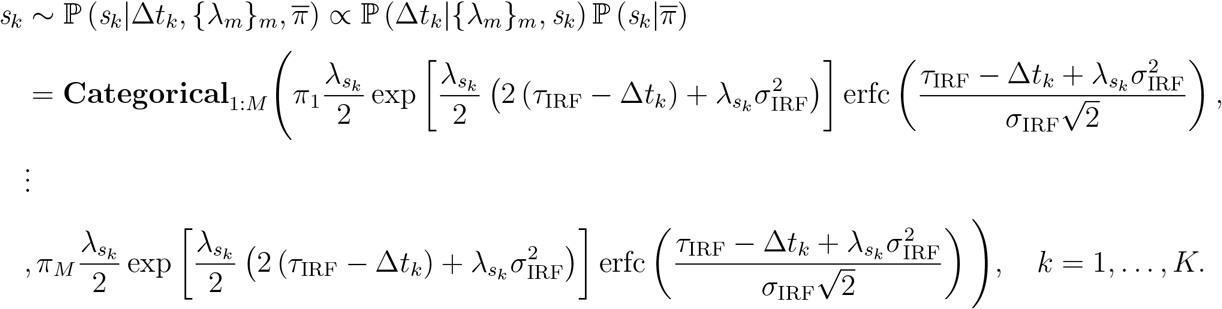

#### Sampling the inverse lifetimes {λ_*m*_}_*m*_

To sample λ_*m*_, we sample from the corresponding conditional probability distribution 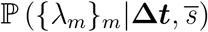 as follows

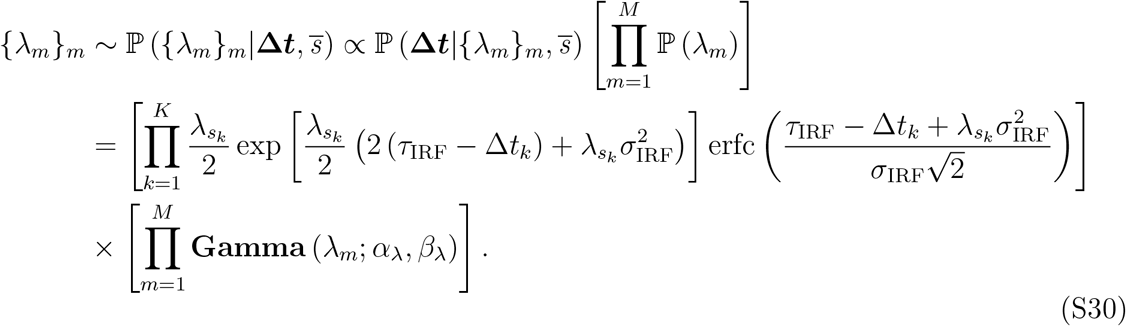

Since, there is no closed form to sample {λ_*m*_}_*m*_, we sample it using the Metropolis algorithm with the proposal

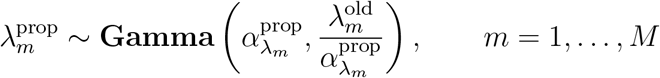

where, the 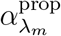 is the parameter of the proposal distributions for the inverse lifetime. Then, the acceptance ratio is equal to

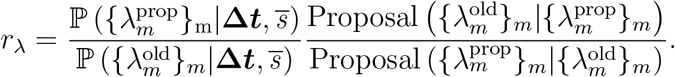

Also, to avoid numerical underflow, we work with the logarithm of the acceptance ratio

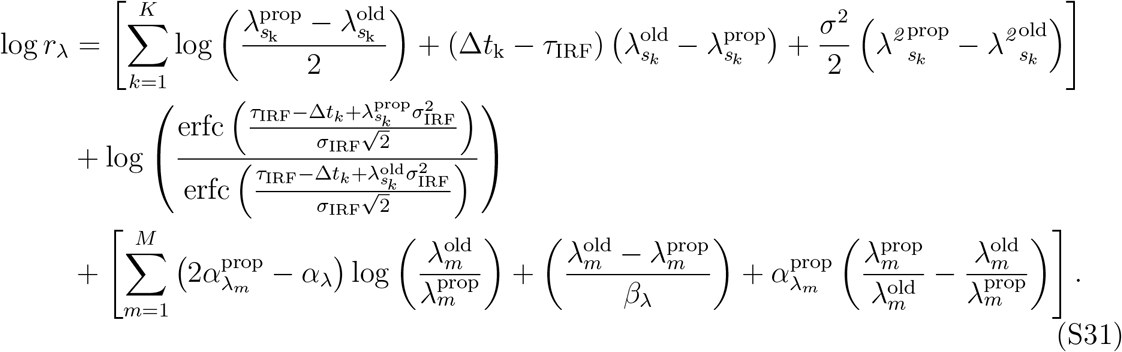

So, at the end we will accept or reject the proposal if

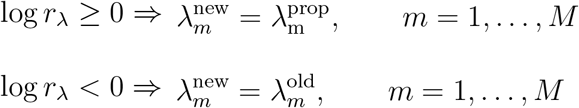

##### Label switching correction of the molecular lifetimes

Label switching is a well-known feature of BNPs.^127^ It arises when we are exploring complex posterior distributions by MCMC algorithms and the likelihood of the model is invariant to the relabelling of mixture components.^128^ The issue of label switching appears because the likelihood is invariant under permutation of the indices. Under symmetric priors, the posteriors also reflects the likelihood’s invariance with respect to index permutation. As a result, in any MCMC algorithm, labels of the components can permute multiple times between iterations of the sampler.^129,130^ Concretely, here, due to exchangeability of the molecular lifetimes, at any iteration (*i*) of the Gibbs sampling scheme, the corresponding lifetime of the species *m* might switch with the molecule’s lifetime of the species *m*′. This label switching does not affect the joint posterior over all lifetimes.

To undo such label switching, at any iteration of the Gibbs sampling we compare the sampled lifetimes 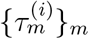 and their weights 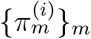 with a fixed set of lifetimes 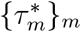 and weights 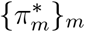. Based on the distances of the lifetimes at iteration (*i*) from the fixed set of lifetimes, which we chose, we correct for label switching. The simple choice for this distance can be the distance between the lifetimes, but, since label switching happens in the sampled lifetimes, and subsequently the weights of each molecular lifetime, the particular distance we use incorporates the emission probability and the weights of each molecular lifetime

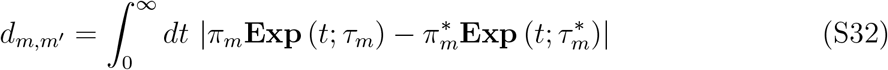

and we solve the assignment problem is minimizing this distance over the species 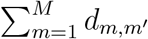. This problem and its computation can be done efficiently by applying the Hungarian algorithm.^131–133^

**Table S1:**
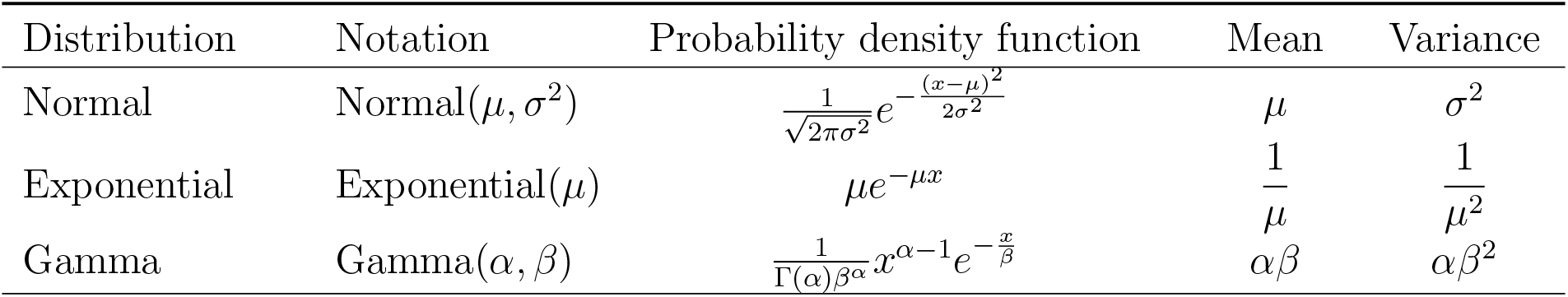
Probability distributions used and their densities. Here, the corresponding random variables are denoted by *x*.

**Table S2:** Here, we list point estimates of our analyses for synthetic data, which we obtain from the marginal posterior probability distributions *p*(*τ*|**Δ*t***). Estimates are listed according to figure.

| | $\tau$ | |
| --- | --- | --- |
|  | mean | std |
|  | ns | ns |
| Fig. 2C | 0.51 , 2.19, 10.51 | 0.14 , 1.42 , 6.45 |
| Fig. 2D | 0.52 , 2.36 , 13. 01 | 0.26 , 1.65 , 12.59 |
| Fig. 2E | 0.52 , 2.51 , 9.74 | 0.31 , 2.33 , 15.74 |
| Fig. 2F | 0.51 , 2.10 , 11.06 | 0.32 , 0.65 , 6.71 |
| Fig. 3B1 | 1.17 | 0.29 |
| Fig. 3B2 | 1.03 | 0.23 |
| Fig. 3B3 | 1.04 | 0.05 |
| Fig. 3B4 | 1.01 | 0.03 |
| Fig. 4B1 | 0.82 , 8.88 | 0.41 , 10.31 |
| Fig. 4B2 | 1.10 , 10.37 | 0.33 , 6.31 |
| Fig. 4B3 | 1.07 , 10.08 | 0.15 , 4.98 |
| Fig. 4B4 | 1.01 , 10.1 | 0.05 , 5.23 |
| Fig. 5A | 0.95 , 9.21 | 0.21 , 8.91 |
| Fig. 5B | 1.10 , 10.13 | 0.35 , 7.11 |
| Fig. 5C | 1.07 , 10.08 | 0.15 , 10.18 |
| Fig. 6A | 1.05 , 10.12 | 0.14 , 3.84 |
| Fig. 6B | 1.10 , 5.11 | 0.25 , 3.11 |
| Fig. 6C | 0.87 , 2.18 | 0.98 , 2.06 |
| Fig. 6D | 1.13 , 1.48 | 0.26 , 0.68 |
| Fig. S1A | 0.85 | 0.31 |
| Fig. S1B | 1.03 | 0.39 |
| Fig. S1C | 0.99 | 0.48 |
| Fig. S1D | 1.01 | 0.11 |
| Fig. S4 | 1.01 , 4.10 , 10.06 | 0.12 , 0.35 , 5.21 |
| Fig. S5 | 0.51 , 1.97 , 6.16 , 12.25 | 0.14 , 0.55 , 3.41 , 7.43 |

**Table S3:** Here, we list point estimates of our analyses for experimental data, which we obtain from the marginal posterior probability distributions *p*(*τ*|**Δ*t***). Estimates are listed according to figure.

| | $\tau$ | |
| --- | --- | --- |
|  | mean | std |
|  | ns | ns |
| Fig. 7A1 | 3.14 | 2.49 |
| Fig. 7B1 | 3.84 | 1.84 |
| Fig. 7C1 | 3.85 | 0.37 |
| Fig. 8A1 | 1.44 , 3.39 | 1.14 , 1.52 |
| Fig. 8B1 | 1.42 , 3.86 | 0.46 , 1.05 |
| Fig. 8C1 | 1.41 , 3.71 | 0.30 , 1.10 |
| Fig. 9A1 | 1.44 , 3.42 | 0.48 , 1.62 |
| Fig. 9B1 | 1.42 , 3.91 | 0.39 , 1.24 |
| Fig. 9C1 | 1.37 , 3.75 | 1.12 , 1.15 |
| Fig. S7 | 0.21 , 1.37 , 2.06 , 3.89 | 0.25 , 0.72 , 1.41 , 2.44 |

## Graphical TOC Entry

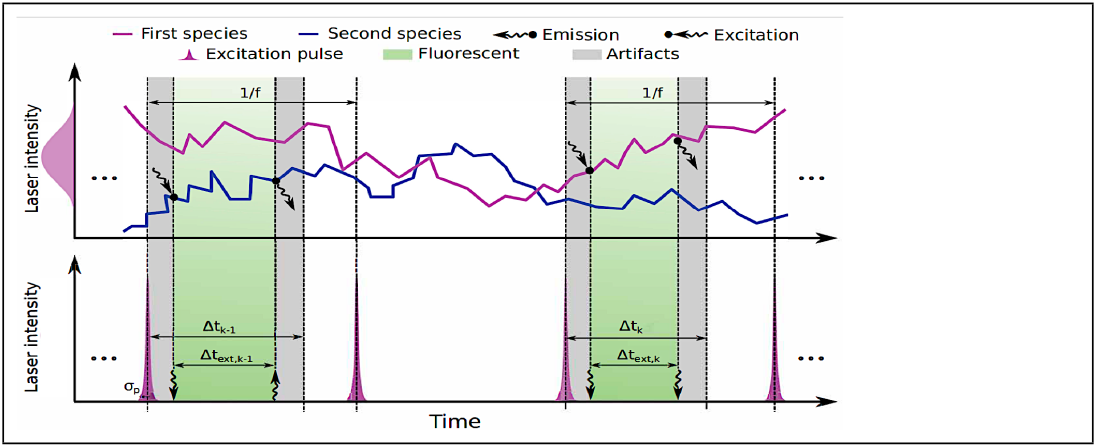

